# Distinct spatial organisation of Rho and RNA Polymerase in *Salmonella* cells

**DOI:** 10.64898/2026.05.02.722398

**Authors:** Lionello Bossi, Romain Le Bars, Johnathan C. Black, Julia Buggiani, Caroline Clerté, Thuy Duong Do, Emmanuel Margeat, Marc Boudvillain, Nara Figueroa-Bossi

## Abstract

Rho is a conserved, ATP-dependent RNA translocase that terminates transcription at hundreds of sites across bacterial genomes. Although the molecular mechanism of Rho-dependent termination is well characterized, its spatial interplay with RNA polymerase (RNAP) within bacterial cells remains elusive. To address this question, we constructed intragenic, in-frame fusions inserting mCherry or sfGFP 48 amino acid residues downstream of the N-terminus of Rho in *Salmonella*. Strikingly, mCherry - but not sfGFP - renders the first 48 residues of Rho dispensable. RhoΔ48::mCherry is viable in single copy and exhibits wild-type termination activity in vitro, whereas the full-length Rho::sfGFP fusion, although viable, slows growth and shows strongly reduced activity. Structured illumination microscopy (SIM) revealed that, despite these functional differences, both constructs exhibit similar localisation patterns relative to fluorescently tagged RNAP in single cells. During exponential growth, both Rho and RNAP form discrete clusters, but with markedly distinct spatial organisations: RNAP clusters associate with the nucleoid, whereas Rho is distributed throughout the cell body. This spatial partitioning persists in stationary phase, where RNAP becomes diffusely associated with a compacted nucleoid while Rho accumulates at the cell periphery. The widespread distribution of Rho at cytoplasmic locations is unexpected and suggests participation in cellular functions beyond its canonical role in transcription termination.

## INTRODUCTION

Bacterial transcription terminates through two alternative pathways [1]. Intrinsic (Rho-independent) termination requires no auxiliary factors and is dependent for the most part on the formation of a stable secondary structure in the nascent RNA. The second pathway depends on the conserved, ATP-driven termination factor Rho. Although intrinsic terminators account for a large proportion of transcriptional endpoints, Rho acts at hundreds of sites and plays a central role in genome surveillance, most notably by suppressing pervasive antisense transcription [2–5]. Rho is a ring-shaped homohexameric ATPase with multiple RNA-binding domains (reviewed in [6]). In *Escherichia. coli*, each Rho subunit consists of 419 amino acids (∼47 kDa), 417 of which are identical in *Salmonella enterica* (99.5% identity). The amino-terminal domain (NTD), comprising roughly the first 130 residues, forms the RNA-binding face of the ring and contains two subdomains: the N-terminal helix bundle (NHB) and the primary RNA-binding site (PBS). The remainder of the protomer forms the C-terminal domain (CTD), which houses the ATP-hydrolysis pocket as well as the secondary RNA-binding site required for translocation. Two mechanistic frameworks have been proposed to explain how Rho engages the elongation complex (EC). In the classical “RNA-centric” or “catch-up” model, Rho first binds unstructured, C-rich rut (Rho-utilization) sites on the nascent RNA and then translocates along the transcript until it reaches RNA polymerase (RNAP), triggering termination by destabilizing the RNA:DNA hybrid or by displacing RNAP’s catalytic center from the 3ʹ end of the RNA [7–11]. The alternative “stand-by” model posits that Rho associates with RNAP early, possibly via NusG or NusA, and scans the emerging transcript directly at the exit channel. Upon encountering a rut site, Rho’s translocase activity induces a conformational change that inactivates RNAP and terminates transcription [12–14]. The two models are not necessarily mutually exclusive, but represent termination pathways that coexist, each optimized for a specific context or condition [15].

The *rho* gene is found throughout the bacterial kingdom, with the notable exception of Cyanobacteria, Tenericutes, and a significant fraction of Firmicutes [16, 17]. In Actinobacteria, Bacteroidetes, and several Firmicutes, Rho contains a large variable insert located at the boundary between the NHB and PBS subdomains [16, 17]. This N-terminal insertion domain (NID) is generally intrinsically disordered and often contains prion-like sequence features. In the human gut commensal *Bacteroides thetaiotaomicron*, the NID was shown to promote liquid-liquid phase separation (LLPS) and the formation of biomolecular condensates that stimulate termination activity [18]. Stimulation of Rho activity by the NID was also reported in *Clostridioides difficile*, but was attributed to a positive influence on hexamerization [19]. In contrast, in *Clostridium botulinum*, the NID drives the assembly of amyloid structures that inactivate Rho [20, 21]. A different type of aggregation, also inhibitory, has recently been reported for *E. coli* Rho. The Psu protein of satellite phage P4, as well as the nucleotides ADP and ppGpp, were shown to induce the formation of higher-order, inactive Rho oligomers [22, 23]. In some instances, such “hyper-oligomerization” proved to be reversible, suggesting that it could constitute a mechanism for temporarily inhibiting Rho activity in response to stress [23]. This would permit a more rapidly adjustable response than the long-known classical feedback mechanism through which Rho down-regulates its own synthesis by attenuating transcription in the leader region of its gene [24, 25].

The potential for Rho to form higher-order structures in vivo renews interest in exploring its spatial relationship with RNA polymerase, an issue made particularly compelling by the demonstration that RNAP itself partitions into subcellular compartments through LLPS in exponentially growing *E. coli* [26]. In the present study, we investigated this relationship by constructing *Salmonella* strains in which the chromosomal *rho* and *rpoC* genes were fused to either *sfGFP* or *mCherry* in the reciprocal combinations. In the Rho constructs, the fluorophore-encoding sequence was inserted intragenically at the NID position, after we found that sfGFP fusions at either the amino (N)- or carboxyl (C)-terminal were inviable in single copy. During these constructions, we made the striking observation that, when inserted at the NID position, mCherry, but not sfGFP, renders the first 48 amino acid (aa) residues of Rho (comprising the entire NHB subdomain) dispensable. The bulk of this study was carried out using two strains: one in which the mCherry construct (*rhoΔ48*::*mCherry*) is combined with a C-terminal sfGFP fusion to *rpoC* (*rpoC-sfGFP*), and another carrying the sfGFP insertion in *rho* (*rho*::*sfGFP*) together with a C-terminal mCherry fusion to *rpoC* (*rpoC-mCherry*). Structured illumination microscopy (SIM) analysis of the dual-labelled strains yielded closely comparable fluorescence patterns regardless of fluorophore arrangement, despite differences in the termination activities of RhoΔ48::mCherry and Rho::sfGFP. We found that both Rho and RNAP display clustering patterns that, somewhat surprisingly, show only limited spatial overlap. Spatial partitioning is particularly evident in stationary phase, where RNAP is mostly associated with the bacterial nucleoid, while Rho accumulates at the cell periphery and poles.

## RESULTS

### Intragenic sfGFP and mCherry insertions at the NID position yield viable single-copy chimeras

We began this work by attempting to construct C-terminal sfGFP fusions to the *rho* and *rpoC* genes in the *Salmonella* chromosome using λ Red–mediated recombineering [27]. While the *rpoC* construct was readily obtained — a result subsequently reproduced with an analogous mCherry fusion — the construction of the *rho* fusion proved problematic. Candidate recombinants were rare and, upon characterization, were found to carry both the desired fusion (as confirmed by DNA sequencing) and a wild-type copy of *rho*. We concluded that these clones originated from cells carrying a tandem duplication of the chromosomal region encompassing the *rho* gene, suggesting that the *rho-sfGFP* fusion is inviable in single copy. It is noteworthy that *rho* is located near the origin of replication (*oriC*), a region prone to duplications due to unequal crossover events between sister chromatids [28]. To confirm this conclusion and obtain a viable fusion, we adopted an unbiased strategy in which fusions were first constructed in a strain carrying a second copy of *rho* elsewhere in the chromosome and subsequently tested for transferability to the haploid background (see Supplementary Information for details). This approach allowed construction of N-terminal and C-terminal fusions, as well as an intragenic fusion with the *sfGFP* ORF inserted between codons 46 and 49 (removing codons 47 and 48), corresponding to the NID position in several bacterial species. Only the intragenic fusion could be transferred to a wild-type background (Fig. S1), establishing this configuration as the sole permissive one for growth in single copy. Based on these results, a second construct with the mCherry ORF inserted at the same NID position was also generated (Fig. 1A). Western blot analysis confirmed that the Rho proteins from strains carrying the single-copy fusions displayed the gel mobility shifts predicted for the fluorophore addition (Fig. 1B).

**Figure. 1.**
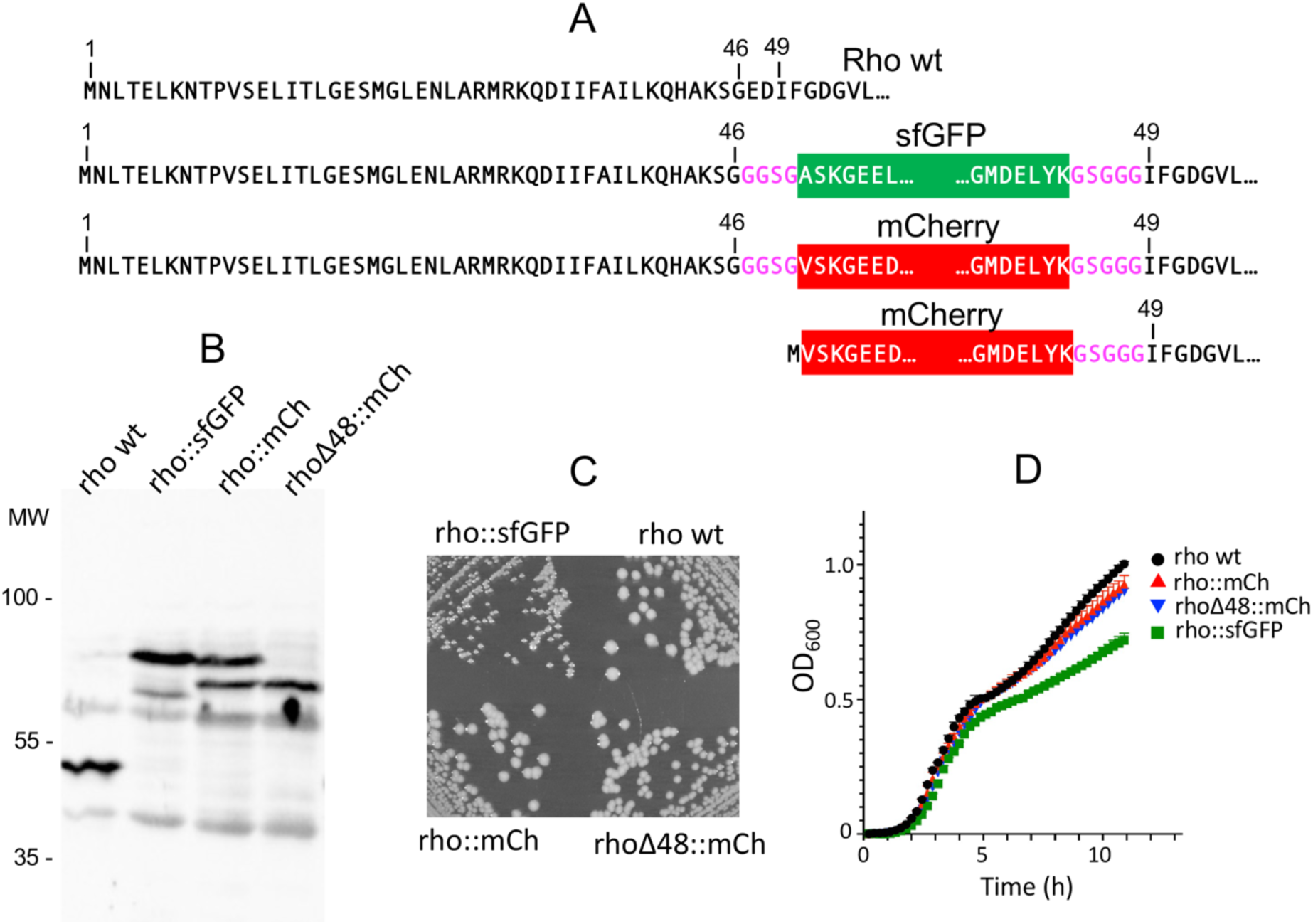
Construc0on and characteriza0on of intragenic fluorescent protein fusions to *rho*. **A**. Amino-acid sequences of the constructs. Black le7ering denotes Rho residues; purple denotes linker sequences. Fluorophore sequences are shown in white le7ering and boxed. From top to bo7om: alleles *rho*::*sfGFP*, *rho*::*mCherry*, and *rho*Δ48::*mCherry*. **B.** Western blot analysis of strains carrying the *rho* fusions. Strains were grown in liquid culture at 37 °C to an OD_600_ of 1, aOer which cells were harvested by centrifugation and pellets processed for Western blot analysis using anQ-Rho polyclonal antibodies, as described in Experimental Procedures. From leO to right: strains MA3409 (rho^wt^), MA12189 (*rho*::*sfGFP*), MA12867 (*rho*::*mCherry*) and MA14978 (*rho*Δ48::*mCherry*). **C**. Colonies from the four strains in B on LB plates. **D.** Growth rates of the same strains in liquid LB measured using a Tecan Infinite 200 Pro plate reader. mCherry is abbreviated mCh in panels **B**-**D**.

### Intragenic mCherry, but not sfGFP, renders Rho’s NHB subdomain dispensable

During sequence verification of isolates obtained from the *rho*::*mCherry* construct, we identified a clone carrying a single base-pair deletion near the upstream boundary of the *mCherry*-encoding sequence. This mutation, which was absent from other isolates and therefore likely introduced during the recombineering step, is predicted to cause a translational frameshift leading to premature termination at the very beginning of the *mCherry* sequence (Fig. S2). This was surprising, as the frameshift would be expected to prevent translation of the downstream *rho* coding sequence. Inspection of the mCherry sequence revealed the presence of an in-frame AUG codon preceded by a putative Shine–Dalgarno motif a short distance from the fusion junction, suggesting a potential site for translational reinitiation (Fig. S2). The functionality of such an internal translation initiation site within the *mCherry* mRNA has recently been experimentally demonstrated [29]. The obligate inference was that the first 48 codons of *rho* — comprising the entire NHB subdomain — are not essential for the viability of the *rho*::*mCherry* fusion. To test this directly, a new *rho*::*mCherry* construct lacking codons 1–48 was generated (Fig. 1A), alongside equivalent deletions in the *rho*::*sfGFP* fusion and in the wild-type *rho* gene. Only the *mCherry* fusion was transferable into a haploid background, demonstrating that mCherry, but not sfGFP, can functionally replace the NHB subdomain, which remains essential in wild-type Rho. Hence, mCherry must possess specific features that allow it to complement the NHB deletion. As expected, the Δ48 allele increased the electrophoretic mobility of the fusion protein in the Western blot (Fig. 1B). The presence of a band co-migrating with this faster species in the blot profile of *rho::mCherry* (Fig. 1B, lane 3) indicates that the internal initiation site is also active in the full-length construct. Notably, while the strain carrying *rho*::*sfGFP* exhibits a slower growth rate, particularly on solid media (Fig. 1C), the isolate carrying *rho*Δ48::*mCherry* grows at a rate comparable to wild-type (Fig. 1C,D).

### RhoΔ48::mCherry retains high transcription termination activity

To further characterize the fluorophore-tagged Rho variants, RhoΔ48::mCherry and Rho::sfGFP were purified and assayed for their ability to promote transcription termination at the bacteriophage λ tR1 terminator in vitro (Fig. 2A). Under standard reaction conditions (100 mM KCl), wild-type Rho promotes termination at multiple positions within the *tR1* region. Addition of NusG stimulates Rho activity, increasing the proportion of transcripts terminated at promoter-proximal sites. As shown in Fig. 2B, RhoΔ48::mCherry exhibits termination activity comparable to that of wild-type Rho. In contrast, no detectable termination products are observed with Rho::sfGFP under these conditions. Lowering the KCl concentration to 50 mM, a more permissive condition for termination, allows Rho::sfGFP to show limited activity in the presence of NusG (Fig. 2C). Finally, replacing KCl with potassium glutamate (KGlu), which further stimulates Rho-dependent termination [30], results in detectable termination products for both chimeric proteins, in both the presence and absence of NusG (Fig. 2D). In 50 mM KCl, Rho::sfGFP also exhibits RNA-dependent ATP hydrolysis activity when provided with an excess of poly(rC), the most active RNA substrate for wild-type Rho [31] (Fig. 2E,F). Overall, however, Rho::sfGFP activity appears markedly reduced, raising the question of how the cell can tolerate such functional defect. Measurements of fluorescence levels in strains carrying the different fusions indicate that the answer lies in the regulation of the *rho* gene. Specifically, the strain harboring the *rho::sfGFP* allele compensates for reduced Rho activity by overexpressing the gene, with expression reaching ∼two-fold higher than *rpoC*. In contrast, the *rhoΔ48::mCherry* allele is expressed at only ∼50% of the *rpoC* level (Fig. 3).

**Figure 2.**
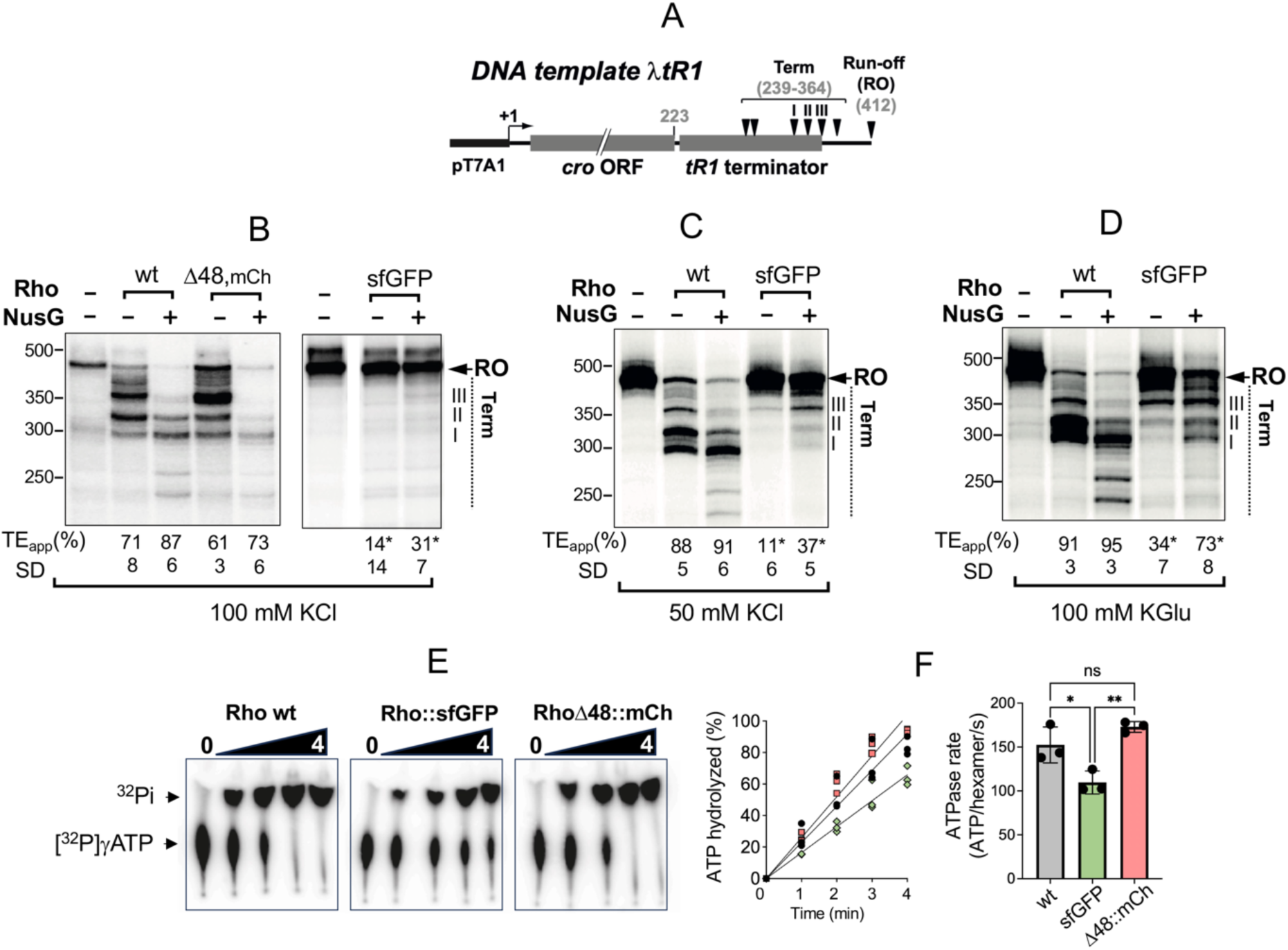
In vitro activity of fluorescent Rho chimeras. **A**. Schematic representation of the DNA template used for in vitro transcription. The template spans the phage λ *tR1*terminator region, which contains multiple sites at which transcription is halted in the presence of Rho, with or without NusG. **B**-**D**. RNA profiles obtained in the presence of 100 mM KCl (**B**), 50 mM KCl (**C**), and 100 mM potassium glutamate (KGlu) (**D**). Quantification values are from 2-6 independent experiments, except for those marked with an asterisk, which originate from double blind quantification of the same gel with no experimental replicate available. Differences in overall band intensities between lanes reflect transcript-length–dependent incorporation of α-[³²P]-UTP and minor variations in reaction loading. Therefore, termination efficiencies were calculated from normalized band intensities rather than raw signal intensity comparisons between lanes. **E**–**F**. Comparison of ATPase activity of wild-type Rho and the fluorescent chimera. (**E**) Representative thin-layer chromatography (TLC) autoradiograms showing the separation of γ-^32^P-labelled ATP and hydrolysis products over the indicated incubation times (minutes). (**F**, left) Time course of ATP hydrolysis plotted as the percentage of ATP hydrolyzed over time. Circles, squares and diamonds specify Rho wt, RhoΔ48::mCherry and Rho::sfGFP, respectively; data points represent three independent experiments. (**F**, right) Bar graph showing the corresponding ATPase rates, calculated from the initial slopes of the hydrolysis curves and expressed as ATP molecules hydrolyzed per hexamer per second (ATP/hexamer/s). Statistical significance was determined by one-way ANOVA with Tukey’s multiple comparisons test (ns, P> 0.05; *, P ≤ 0.05, ** P≤ 0.01).

**Figure 3.**
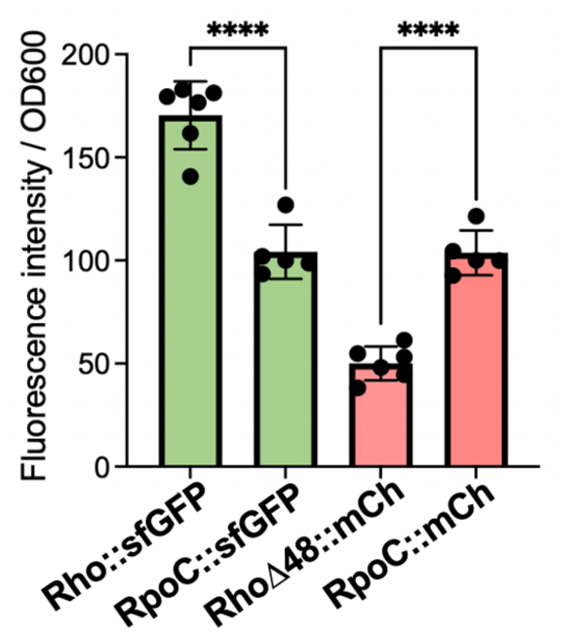
Relative fluorescence of Rho and RpoC fusions. Exponentially growing cultures from *Salmonella* strains MA12147 (*rpoC*-*mCherry*), MA12451 (*rho*::*sfGFP*), MA12859 (*rpoC*-*sfGFP*) and MA14978 (*rho*Δ48::*mCherry*) were measured for total fluorescence using a Tecan Infinite 200 Pro plate reader. Fluorescence values were normalized to the cell density (OD600) and to the mean fluorescence of the corresponding RpoC fusion (set to 100). Data represent the mean ± SD of three independent experiments. Statistical significance was determined by one-way ANOVA with Šidák’s multiple comparisons test (****, P ≤ 0.0001).

Interestingly, AlphaFold3 structural predictions [32] offer a mechanistic explanation for the markedly different activities of the fluorescent Rho chimeras. Despite their structural similarity, mCherry and sfGFP have distinct electrostatic surface potentials (Fig. S3), which are predicted to influence their preferred orientation relative to the Rho hexamer. In the mCherry fusion, the fluorophore β-barrel is predicted to dock preferentially on the side of the Rho ring, leaving the RNA-binding surface on the top of the ring fully accessible (Fig. 4A,B, Fig. S4A). This lateral configuration is maintained in the Δ48 derivative, where a slt of the barrel axis brings the connecting sequence closer to the top surface (Fig. 4C). This connector may contribute to the formation of a positively charged, crown-like surface on the Rho hexamer (Fig. S3) — a feature thought to be critical for RNA capture — which might explain mCherry’s ability to complement the lack of the NHB domain. In contrast, in the Rho::sfGFP chimera, the sfGFP barrel is predicted to lie flat atop the hexameric ring, a configuration likely to interfere with RNA binding (Fig.s 4D,E, Fig. S4B). In this context, Rho activity may become strictly dependent on the presence of the NHB, providing a plausible explanation for the lethality of Δ48 in the sfGFP fusion context.

**Figure 4.**
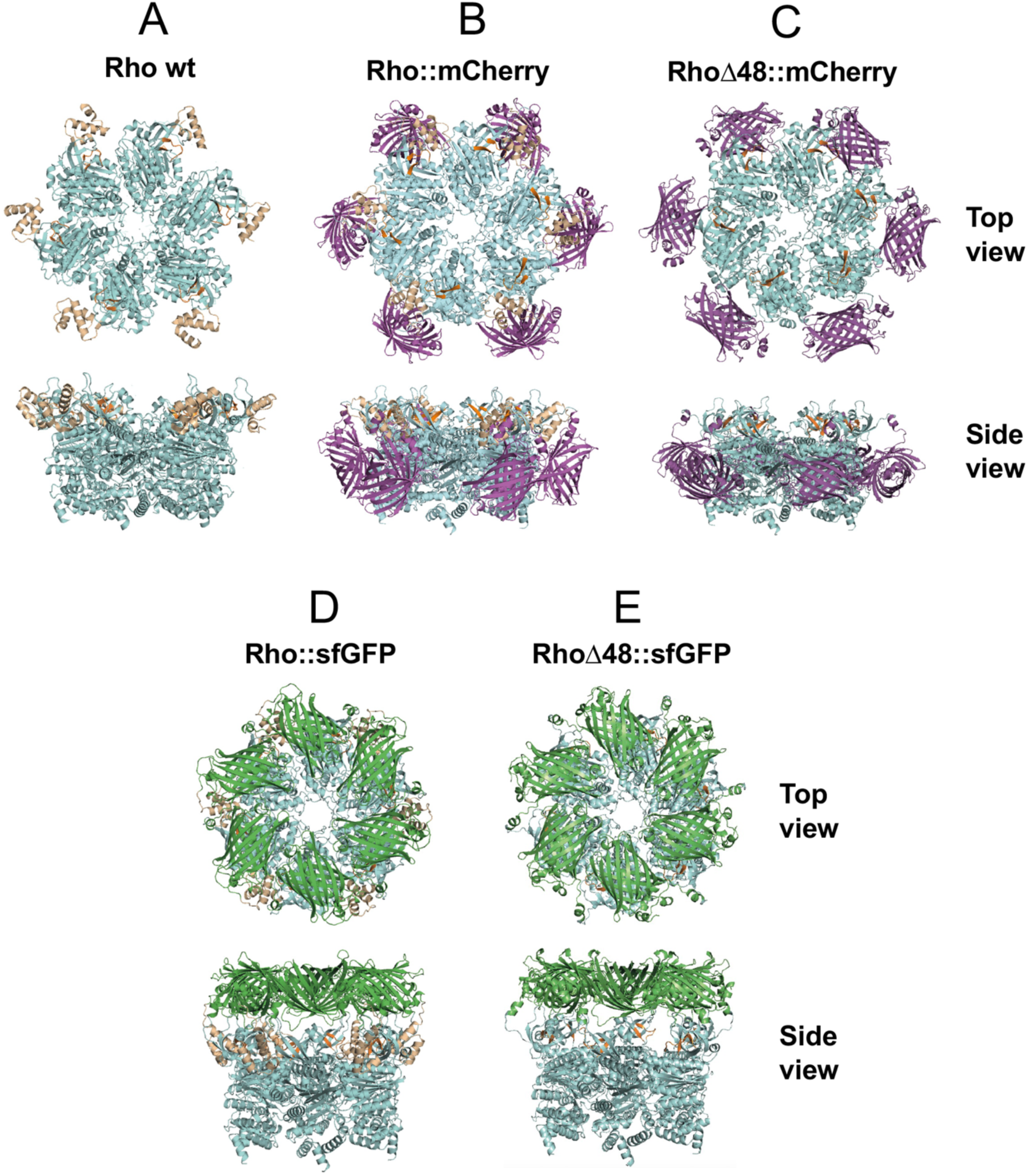
Predicted structures of Rho fusion proteins. Models of the RhoΔ48::mCherry and Rho::sfGFP chimeras were generated using AlphaFold 3 via the AlphaFold Server (alphafoldserver.com), with a 6-mer constraint applied. For comparison, the crystal structure of wild-type (WT) Rho (PDB: 3ICE) is also shown. AlphaFold per-atom confidence (pLDDT) and Electrostatic potential maps are shown in Figures S3 and S4, respectively. The mCherry and sfGFP insertions are colored magenta and green, respectively, while the Rho C-terminal domain (CTD) is shown in cyan. Residues forming the PBS pockets are highlighted in orange, and residues 1–48 (as in WT Rho) are depicted in wheat. All structural images were prepared using PyMOL v2.6.

To further examine possible steric effects, we performed additional AlphaFold simulations including the lambda tR1 rut site as a model RNA ligand. In these models, RNA engages the primary binding site in all constructs, but the sfGFP fusion places the fluorescent moiety closer to the site than mCherry, suggesting a stronger steric interference rather than complete exclusion of RNA binding (Fig. S5).

### Rho and RNA Polymerase form largely non-overlapping clusters in single cells

Dual-color structured illumination microscopy (SIM) was performed on strains expressing either of the two Rho fusions in combination with a C-terminally fluorescently labelled RpoC. In exponentially growing cells, both Rho::sfGFP and RhoΔ48::mCherry show diffuse, patchy fluorescence patterns distributed throughout the cell body, including regions proximal to the cell periphery (Fig. 5A,C). Only a limited subset of these patches appears to overlap with RNAP and/or the nucleoid, in contrast to RNAP foci, which are more closely associated with nucleoid regions. The notable similarity in the spatial distributions of the two Rho alleles — despite their marked differences in termination activity — suggests that the observed clustering patterns reflect intrinsic features of Rho cellular organisation rather than fluorophore-specific effects. Rho’s segregation from RNAP is also evident in stationary phase. Under these conditions, RNAP clusters largely dissipate and the protein becomes diffusely associated with a more compacted nucleoid, whereas Rho accumulates toward the cell periphery and the poles (Fig. 5B,D).

**Figure 5.**
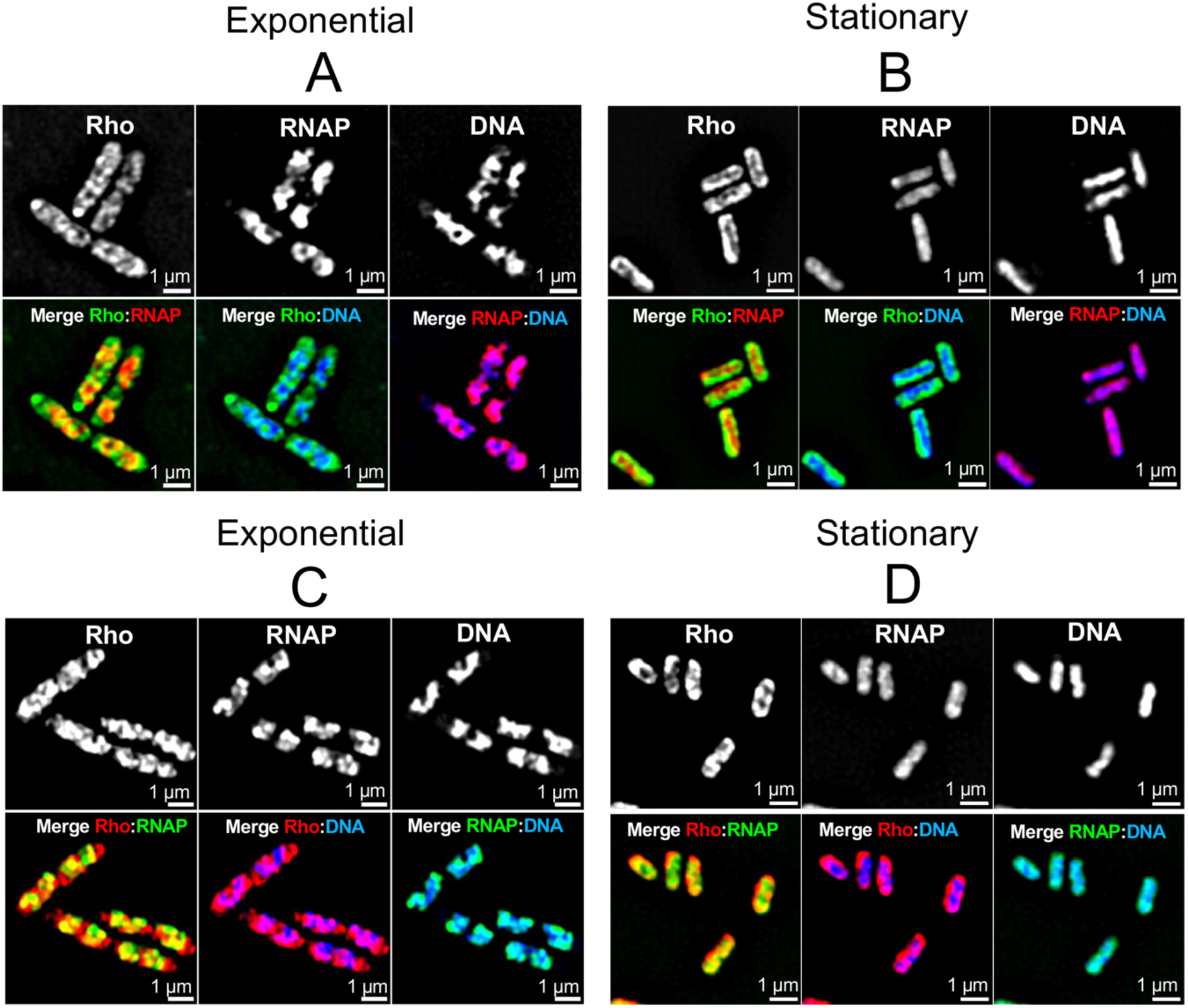
Dual-color SIM imaging of sfGFP- and mCherry-tagged Rho and RNAP and DAPI-stained DNA in single cells. Bacteria were grown to an OD600 of 0.2 (exponential phase) or to an OD600 of 2 (stationary phase), fixed with paraformaldehyde, and processed for structured illumination microscopy (SIM) as described in Experimental Procedures. Images shown are representative of the observed localisation patterns across multiple cells and independent experiments. **A**,**B**. Strain MA12189 (*rho*::*sfGFP rpoC*-*mCherry*) in exponential phase (**A**) and stationary phase (**B**). **C**,**D.** Strain MA14978 (*rho*Δ48::*mCherry rpoC*-*sfGFP*) in exponential phase (**C**) and stationary phase (**D**). In exponential phase, Rho and RNAP (RpoC) show patchy fluorescence patterns with only partial spatial overlap. In stationary phase, RNAP patches become more diffuse and remain associated with a compacted nucleoid, whereas Rho accumulates toward the cell periphery. Both reciprocal labelling arrangements yield similar patterns, suggesting that Rho’s spatial organisation is independent of the fluorophore used

The SIM images were analysed using two complementary approaches: an object-based colocalisation algorithm to assess spatial relationships between discrete Rho and RNAP foci, and Manders coefficient analysis to quantify pixel-wise signal overlap across the entire molecular population. To specifically examine cluster relationships, individual fluorescent foci were identified as discrete objects, and spatial association was assessed based on the proximity of their centers of mass in pairwise channel comparisons (Fig. 6). Consistent with the spatial patterns apparent from visual inspection, this approach showed that Rho and RNAP clusters exhibit limited overlap, with 15–30% of clusters colocalizing, independent of fluorophore arrangement, in both exponential and stationary phase.

**Figure 6.**
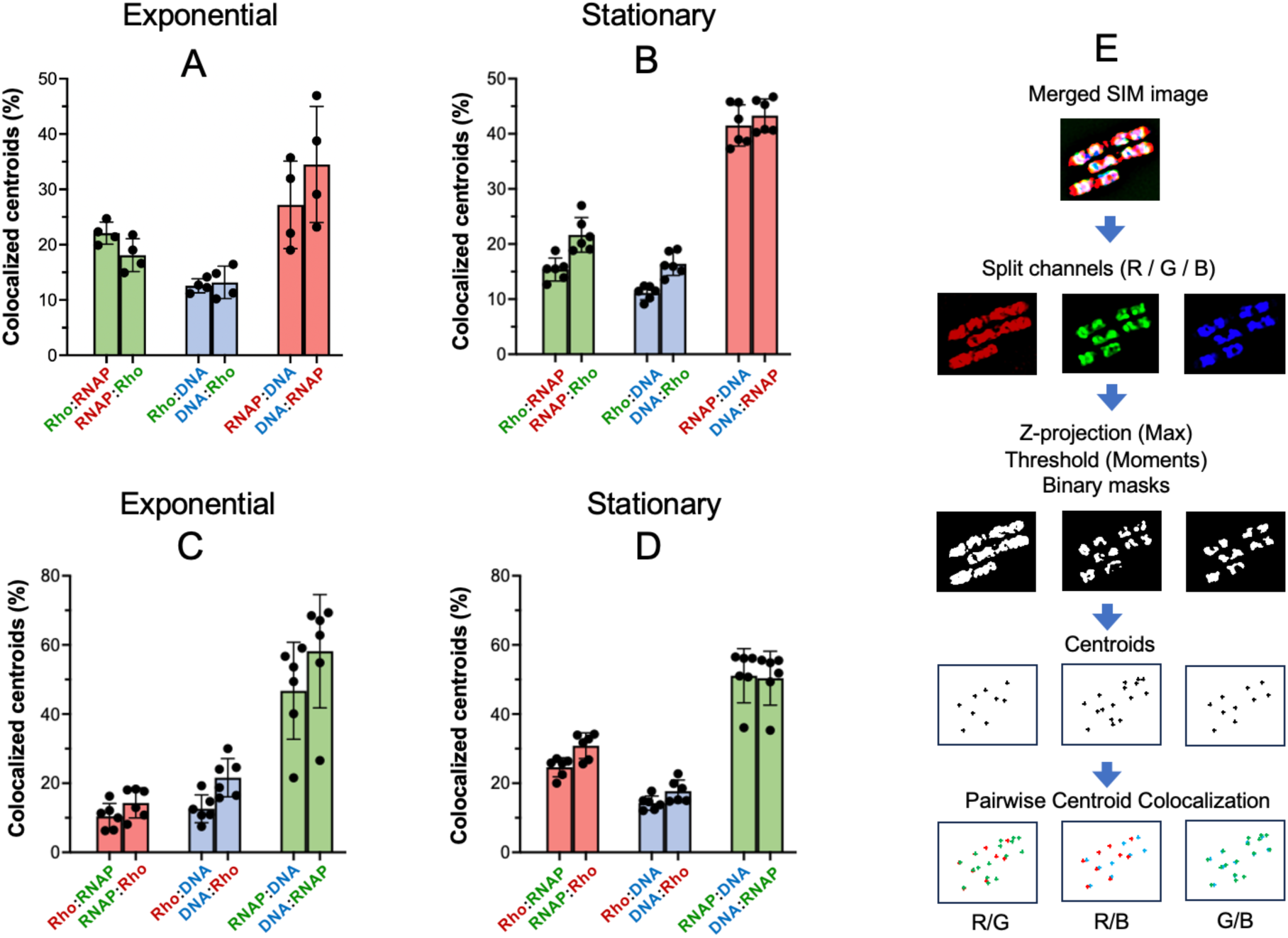
Object-based colocalisation analysis. Z-stacks acquired by structured illumination microscopy (SIM) were processed using SIM2 to reconstruct high-resolution images across three fluorescence channels, followed by maximum-intensity projection. Projected images were thresholded (Moments method) to generate binary masks in which discrete fluorescent patches (referred to here as “clusters”) were defined as individual objects. The centers of mass of these objects were then used for object-based colocalisation analysis using the JACoP plugin in Fiji, which identifies colocalizing objects based on centroid proximity as implemented in the plugin [69]. Each pairwise comparison is represented by two adjacent bars, showing the percentage of centroids from either member of the pair that colocalise with those of the other. **A**,**B**. Strain MA12189 (*rho*::*sfGFP rpoC*-*mCherry*) in exponential phase (**A**) and stationary phase (**B**). **C**,**D**. Strain MA14978 (*rho*Δ48::*mCherry rpoC*-*sfGFP*) in exponential phase (**C**) and stationary phase (**D**). Bars represent mean ± SD from four to six microscopy fields (raw data in Table S4). **E** Schematic representation of the workflow used in **A**–**D**. The procedure is illustrated on a representative subset of cells for clarity.

Manders analysis revealed that in exponential phase roughly 50% of the Rho signal overlaps with RNAP (Fig. 7). The overlap is higher using this method because it captures signal outside discrete cluster areas; nonetheless, it confirms that a substantial fraction of Rho molecules occupies regions lacking detectable RNAP and DNA, consistent with a cytoplasmic pool outside the nucleoid. In contrast, nearly 80% of the RNAP signal colocalises with Rho, indicating that most RNAP molecules are detected in regions enriched in the diffuse, non-clustered pool of Rho. Relative spatial relationships are largely maintained in stationary phase, except for a marked increase in the fraction of Rho colocalizing with RNAP, likely reflecting broadening of the RNAP signal combined with reduced cell volume.

**Figure 7.**
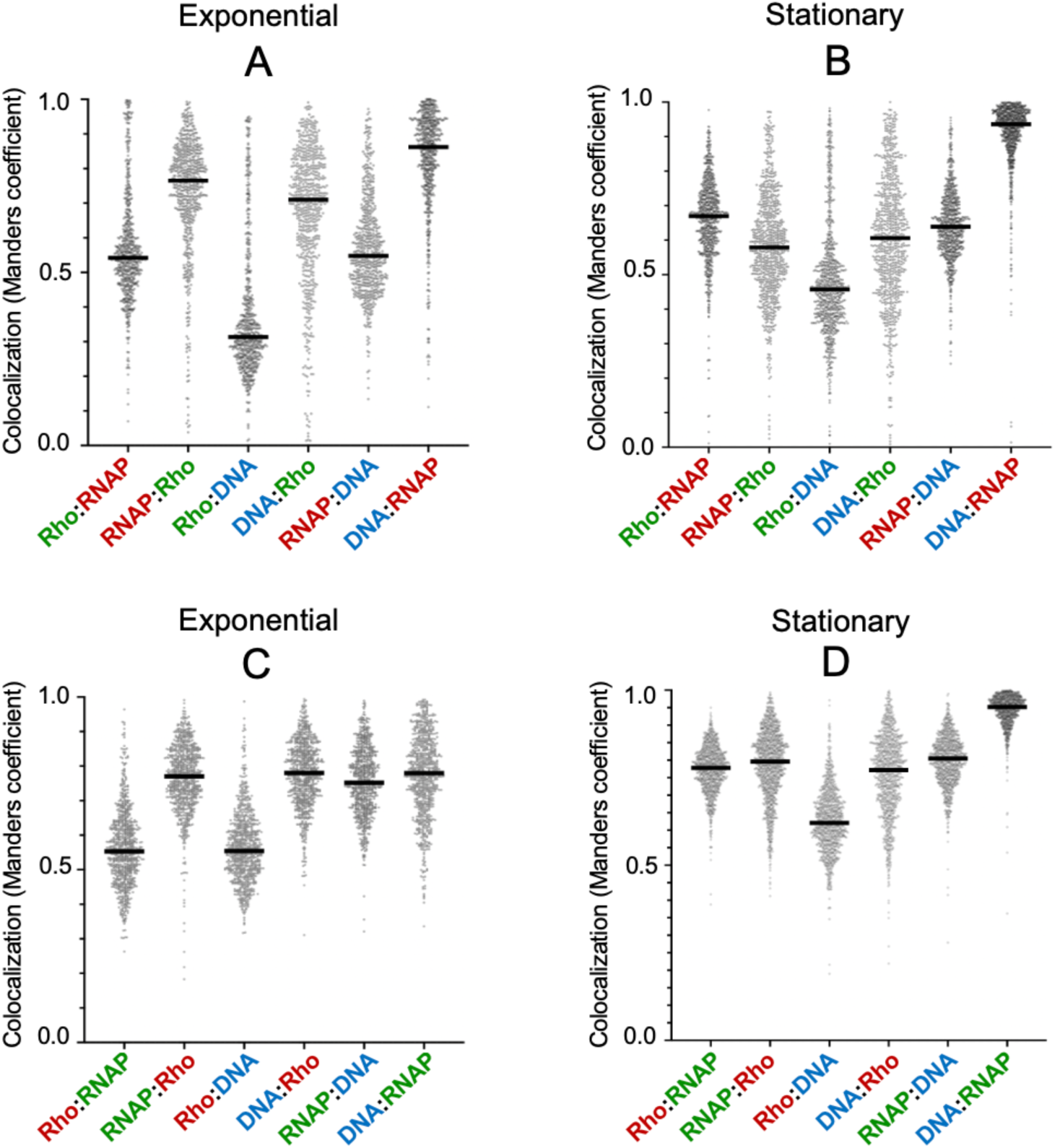
Pixel-based colocalisation analysis. Z-stacks acquired by structured illumination microscopy (SIM) were processed using SIM2 to reconstruct high-resolution images across three fluorescence channels. The resulting datasets were used to generate segmentation masks for individual bacterial cells, as described in Supplementary Experimental Procedures (Fig. S6). Three-dimensional colocalisation was quantified using Manders’ overlap coefficient, with intensity thresholds determined using the Moments method implemented in Fiji (JACoP plug-in) [69]. **A**,**B**. Strain MA12189 (*rho*::*sfGFP rpoC*-*mCherry*) in exponential phase (**A**) and stationary phase (**B**). **C**,**D** Strain MA14978 (*rho*Δ48::*mCherry rpoC*-*sfGFP*) in exponential phase (**C**) and stationary phase (**D**). Black bars indicate median values.

## DISCUSSION

In the present work, we constructed and characterized intragenic fluorescent protein fusions to the Rho factor and assessed their subcellular localisation relative to the C-terminally fluorescently labelled RNAP βʹ subunit, RpoC, in *Salmonella* cells using SIM imaging. Construction of the *rho*::*mCherry* fusion led to the discovery that the mCherry sequence can functionally replace the 48-amino-acid NHB subdomain of Rho. Not only is the RhoΔ48::mCherry chimera viable, but it also supports a normal growth rate in LB under standard conditions and exhibits wild-type levels of transcription termination activity in vitro. In contrast, the NHB subdomain remains essential in the Rho::sfGFP fusion. Although viable in single copy, this fusion slows growth and sharply reduces Rho’s in vitro activity. Despite these functional differences, RhoΔ48::mCherry and Rho::sfGFP show highly similar localisation patterns in single cells. Both constructs appear as patchy foci distributed throughout the cell body, with only limited overlap with RNAP clusters, which are confined to the bacterial nucleoid. While the similarity in the localisation patterns of the two chimeras suggests that they likely reflect intrinsic features of Rho spatial organisation in the cell, we cannot exclude the possibility that the fluorescent tag influences the observed distribution. Nonetheless, our results support the conclusion that normal cell growth does not require detectable enrichment of Rho at the nucleoid under the conditions tested. We also note that previous immunofluorescence analyses of wild-type Rho reported signal both within and outside nucleoid regions [33]. While the resolution of these images is limited, they are broadly consistent with the absence of strong nucleoid-restricted localisation and support the idea that the localisation patterns described here are compatible with native Rho behavior.

A recent study examined the cellular localisation of an mCherry–Rho N-terminal fusion in *Escherichia coli* [34]. The authors reported that Rho was primarily localised toward the cell poles during exponential growth, whereas in stationary phase the protein appeared more uniformly distributed throughout the cell. These observations are somewhat difficult to reconcile, particularly because the cells labelled as “exponential phase” in this publication are significantly smaller than those labelled as “stationary phase”. This is opposite to the well-established relationship between cell size and growth phase (see, for example [35]). If the assignment of growth-phase labels were interchanged, the fluorescence payerns reported in that study would be in general agreement with our results.

RNAP clustering in exponentially growing cells has been described in multiple studies and linked to ribosomal RNA (*rrn*) operon transcription [36–42], which is estimated to account for 80–90% of total transcription during rapid growth [26, 38]. One study provided evidence that these RNAP clusters are biomolecular condensates formed through liquid–liquid phase separation (LLPS) [26]. The authors proposed that RNAP is maintained at high local concentration within these condensates through weak multivalent protein–protein interactions. Notably, they presented evidence for the presence of the antitermination factors NusA and NusB within RNAP condensates and for their contribution to condensate formation and stability. NusA and NusB, together with additional factors, form an antitermination complex that prevents premature Rho-dependent transcription termination within rrn operons [43, 44]. If clustered RNAP within condensates is primarily devoted to *rrn* transcription, the limited overlap between RNAP and Rho clusters observed in our study is therefore not surprising. One can further speculate that the involvement of NusA and NusB in condensate formation may itself contribute to antitermination by limiting the physical access of Rho to RNAP within these structures. On the other hand, exclusion of Rho from RNAP condensates would increase the pool of Rho available for termination elsewhere in the genome. In line with this view, our total fluorescence measurements indicate that in exponentially growing cells the *rho*Δ48::*mCherry* fusion — expected to be transcribed at the same level as wild-type *rho* (see Results) — is expressed at approximately 50% of the level of *rpoC*. This ratio is consistent with absolute proteomic measurements in *E. coli*, which estimate ∼7,000 RpoC subunits and ∼6,000 Rho subunits per cell during exponential growth in LB medium [45]. Taking into account that nearly half of cellular RNAP molecules are engaged in *rrn* transcription [26] and that functional Rho acts as a hexamer, these values suggest that exponentially growing cells contain on the order of ∼1,000 Rho hexamers, compared with ∼3,000–4,000 RNAP molecules engaged in transcription of protein-coding genes. This apparent imbalance — further accentuated by the observation that a substantial fraction of Rho localises away from the nucleoid — raises the question of how Rho efficiently recognizes hundreds of binding sites cotranscriptionally in vivo. One possible explanation is that the subset of Rho clusters that colocalise with the nucleoid locally concentrates the factor, thereby facilitating rapid engagement with nascent transcripts. By contrast, the broader distribution of the remaining clusters suggests that they may support activities other than transcription termination, such as roles in mRNA turnover (see below). As a powerful RNA translocase, Rho could contribute to remodeling of mRNA structures refractory to DEAD-box helicases or to resolving stalled ribosomes. Such RNA unfolding activity could also be directed toward Rho-terminated transcripts, which presumably accumulate during to genome-wide suppression of pervasive transcription [4, 5].

The nature of the Rho clusters remains to be determined. One interesting possibility is that they correspond to the higher-order oligomeric assemblies of Rho recently observed in vitro for *E. coli* Rho [22, 23]. Rho hyper-oligomerization was shown to be induced by ADP and (p)ppGpp through distinct mechanisms and was proposed to serve as strategy to curb Rho activity when it is no longer needed [23]. This model could account for the payerns observed in stationary phase, when (p)ppGpp levels are known to increase and we found Rho to accumulate at the cell periphery and poles. However, it does not readily explain the clusters observed during exponential growth. None of the Rho variants analysed in this study contains large intrinsically disordered domains, which at first glance argues against these clusters representing biomolecular condensates formed by LLPS. Nevertheless, Rho could function as a “client protein”, becoming incorporated into condensates scaffolded by other factors without itself driving phase separation [46]. One potential candidate for such a scaffolding compartment is the bacterial degradosome. In *Caulobacter crescentus*, the degradosome has been shown to form condensates driven by the intrinsically disordered region of RNase E [47] and to colocalise with the nucleoid [48]. Rho associates with the degradosome via interactions with RNase E in *Rhodobacter capsulatus* [49, 50] and has also been identified as a component of a large, cyclic-di-GMP-dependent protein complex containing degradosome components in *Escherichia coli* [51]. Notably, unlike in *Caulobacter*, RNase E and other degradosome components in *E. coli* predominantly associate with the inner membrane [52, 53]. In this context, the Rho clusters described here show a spatial organisation reminiscent of the patchy degradosome distribution reported in *E. coli* [53]. Future imaging experiments combining *rho*Δ48::*mCherry* with sfGFP-tagged degradosome components in *Salmonella* should help determine whether these similarities reflect a physical or functional association between Rho and degradosome assemblies. Overall, our study reveals that Rho forms distinct, dynamically positioned clusters that are largely segregated from RNAP, pointing to a multifaceted in vivo organisation in which nucleoid-associated and cytoplasmic Rho populations may serve complementary roles in transcription regulation and RNA metabolism. These findings lay the groundwork for future studies aimed at dissecting their functional relationships with RNAP and other cellular assemblies.

## MATERIALS & METHODS

### Bacterial strains and culture conditions

Strains used in this work are derived from *Salmonella enterica* serovar Typhimurium strain MA3409, a derivative of strain LT2 [54] cured of the Gifsy-1 prophage [55]. The genotypes of the relevant strains used are listed in Table S1. Bacteria were cultured at 37 °C in liquid media or on media solidified by the addition of 1.5% (w/v) Difco agar. Lysogeny broth (LB) [56] was used as a complex medium. Carbon-free medium (NCE) [57], supplemented with 0.2% (w/v) glucose, was used as minimal medium. When needed, antibiotics (Sigma-Aldrich) were included in growth media at the following final concentrations: chloramphenicol, 10 µg mL⁻¹; kanamycin monosulphate, 50 µg mL⁻¹; sodium ampicillin, 100 µg mL⁻¹; spectinomycin dihydrochloride, 80 µg mL⁻¹; tetracycline hydrochloride, 25 µg mL⁻¹. 5-bromo-4-chloro-3-indolyl-β-D-galactopyranoside (Xgal; Sigma-Aldrich), 40 µg mL⁻¹, was included to monitor *lacZ* expression in bacterial colonies.

### Genetic techniques

Strain construction was performed either by generalized transduction using the high-frequency transducing mutant of phage P22, HT105/1 int-201 [58], or by λ Red recombineering implemented as described in [59], [27], and [60]. DNA oligonucleotides used as primers for PCR amplification were obtained from Sigma-Aldrich. The primer–template combinations used to generate the different recombineering constructs analysed in this work are listed in Table S2, and primer sequences are provided in Table S3. PCR-amplified fragments used for recombineering were generated with high-fidelity Phusion DNA polymerase (New England Biolabs). These fragments were introduced by electroporation (Bio-Rad Gene Pulser) into strains carrying the λ Red helper plasmid pKD46 [59]. Recombinant constructs were verified by colony PCR using Taq DNA polymerase, followed by DNA sequencing (performed by Eurofins-GATC Biotech) For details on the construction of relevant strains see Supplementary Methods. Details on the construction of the relevant strains analysed in this study are provided in the Supplementary Experimental Procedures.

### Western blot analysis

Overnight cultures of the strains to be analysed were diluted 1:100 in 2 mL of LB and grown with shaking (170 rpm) at 37 °C to an OD₆₀₀ of 1. Cells were harvested by centrifugation (3 min at 9,000 × g) at 4 °C. Bacterial pellets were resuspended in 100 µl of 2× Laemmli buffer and boiled for 5 min. The resulting whole-cell lysates were separated by 10% SDS-PAGE. PageRuler^TM^ Plus Prestained Protein ladder (Thermo Scientific) was included as migration marker. Proteins were electro-transferred to a PVDF membrane, which was blocked with TBS containing 0.05% Tween-20 and 5% (w/v) powdered low-fat milk. Membranes were incubated with polyclonal antibodies raised against E. coli Rho protein, followed by goat anti-rabbit HRP-conjugated secondary antibodies (Sigma) for 1 h. After incubation, membranes were washed thoroughly with TBS containing 0.05% Tween-20. Detection was performed using the SuperSignal^TM^ West Pico PLUS Chemiluminescent Substrate (Thermo Scientific) and chemiluminescent signals were imaged on a ChemiDoc Touch system (Bio-Rad).

### Proteins for in vitro assay

*E. coli* RNA polymerase (RNAP) holoenzyme was purchased from New England Biolabs. Plasmid for overexpression of the sfGFP-Rho chimera was obtained by subcloning a PCR amplicon encoding the sfGFP insert (Table S5) between the NdeI and NcoI sites of plasmid pET28b-Rho (kindly provided by Pr. James Berger; Johns Hopkins University, USA). Plasmid pET28b(+)_Delta49mCherryRho for overexpression of the RhoΔ48::mCherry protein was obtained by custom plasmid synthesis (Genscript). All Rho protein constructs carry a N-terminal MGH triplet, which increases overexpression while the starting methionine is spontaneously cleaved post-translationally (Table S5) [61]. The Rho and NusG proteins were prepared as described previously [62] with minor modifications (see Supporting Information).

### In vitro transcription

Linear DNA template encoding phage λ tR1 terminator was prepared by standard PCR, as described previously [63]. Transcription termination experiments were performed as described previously [64], with minor modifications. Briefly, DNA template (0.1 pmol), *E. coli* RNAP holoenzyme (0.4 pmol), Rho derivative (0 or 1.4 pmol hexamer), NusG (0 or 2.8 pmol), and SUPERase•In (0.5 U µL^−1^; ThermoFisher Scientific) were mixed in 18 µL of transcription termination buffer (40 mM Tris-HCl, pH 7.9, 5 mM MgCl_2_, 1,5 mM DTT) supplemented with 50 or 100 mM KCl or 100mM potassium glutamate. Mixtures were incubated for 10 min at 37°C before addition of 2 µL of initiation mix (2 mM ATP, GTP, and CTP, 0.2 mM UTP, 2.5 µCi/µL of ^32^P-αUTP, and 250 µg mL^−1^ rifampicin in transcription buffer). Then, mixtures were incubated for 20 min at 37°C before addition of 4 µL of EDTA (0.5 M), 6 µL of tRNA (0.25 mg mL^−1^), and 80 µL of sodium acetate (0.42 M) to quench enzymatic reactions. Mixtures were extracted with a phenol:chloroform:isoamyl alcohol [25:24:1] mix and precipitated at −20°C with 330 µL of ethanol. Reaction pellets were dissolved in denaturing loading buffer (95% formamide, 5 mM EDTA), and analysed by denaturing 8% PAGE and Typhoon FLA-9500 (GE Healthcare) phosphorimaging.

Apparent termination efficiencies (TE_app_) were determined as previously described [63]. Briefly, the lengths of the major transcript species were deduced from the sequence of the λ tR1 DNA template and the reported positions of the three principal Rho-dependent termination clusters [65]. The lengths of minor termination products (150-300 nt range) were estimated from gel migration using the molecular size calculation function of ImageQuant TL (GE Healthcare), calibrated with a low molecular weight DNA ladder (New England Biolabs). For each transcript species, the number of uridine residues (N_U_) was calculated from its length and the sequence of the λ tR1 template. Because transcripts are internally labelled with α-[³²P]-UTP, longer transcripts incorporate more label and therefore yield higher signal intensities. To correct for this bias, band intensities were normalized by the number of uridines. Specifically, for each transcript, a normalized band intensity (I_norm_) was calculated from the raw (uncorrected) band intensity (I_raw_) as follows: I_norm_ = I_raw_ / N_U_. The apparent termination efficiency (TE_app_) for each lane was then calculated from the normalized intensities of truncated (I_norm_^[Trunc]^) and runoff (I_norm_^[Runoff]^) transcripts: TE_app_ (%) = 100 × (ΣI_norm_^[Trunc]^ − TE_0_ × (I_norm_^[Runoff]^+ ΣI_norm_^[Trunc]^)) / ((1 − TE_0_) × (I_norm_^[Runoff]^+ ΣI_norm_^[Trunc]^)), where ΣI_norm_^[Trunc]^ represents the sum of normalized intensities of all terminated species and TE_0_ represents the background termination level, determined from the control (−Rho) lane as: TE_0_ = ΣI_norm_^[Trunc]^ / (I_norm_^[Runoff]^ + ΣI_norm_^[Trunc]^).

### Measurement of Rho’s ATPase activity

ATP hydrolysis rates were determined using a thin layer chromatography (TLC)-based ^32^P radiometric assay, as described previously [66]. Briefly, Rho derivative (0.5 pmol hexamer) and poly(rC) (200 pmol rC residues) were combined in 18 µL of ATPase buffer (50 mM KCl, 1 mM MgCl_2_, 0.1 mM DTT, 0.1 mM EDTA, 20 mM HEPES, pH 7.9) and incubated for 3 min at 37°C. Then, 2 µL of ATP (10 mM, spiked with ^32^P-γATP) in ATPase buffer were added, and the reactions were further incubated at 37°C. Aliquots were withdrawn at specified time points, quenched with two volumes of 0.5 M EDTA, and spotted onto PEI cellulose TLC plates (Macherey-Nagel). Plates were developed with 0.35 M KH_2_PO_4_ (pH 7.5) to separate ^32^P-γATP from the faster migrating ^32^P product, and then imaged with a phosphor screen and a Typhoon FLA9500 instrument (GE Healthcare). The ^32^P-γATP and ^32^P signals were quantified with ImageQuant TL 8.1 (GE Healthcare). The fraction of ^32^P formed (F) was plotted over time and the data fitted to a linear equation, F = *a* x t. The steady-state ATPase rates (in number of ATP molecules hydrolyzed per Rho hexamer and per second) were derived from the slopes of the curves.

### SIM analyses

#### 1. Bacterial fixation and DNA staining

Bacteria in exponential phase (OD₆₀₀ = 0.20–0.25) or stationary phase (OD₆₀₀ = 2) were fixed by gentle mixing with an equal volume of 4% paraformaldehyde. After incubation for 20 min at 4°C, cells were harvested by centrifugation at 4,000 × g and washed twice with 1× PBS. Pellets were resuspended in GTE buffer containing 1 µg mL⁻¹ 4ʹ,6-diamidino-2-phenylindole (DAPI) and incubated at room temperature for 10 min. Cells were then centrifuged again, washed once with 1× PBS, resuspended in 1× PBS, and used for microscopy.

#### 2. Image acquisition

Bacterial samples (1 µL) were deposited onto a 1% agarose pad and covered with a #1.5H coverslip for imaging. Data were acquired using a Zeiss Elyra 7 Lattice Structured Illumination Microscope equipped with a 63× Plan-Apochromat oil-immersion objective (NA 1.4) and a sCMOS PCO.edge 4.2 camera. Fluorescence was excited using 561 nm (mCherry), 488 nm (GFP), and 405 nm (DAPI) lasers, with emission detected through a quad-band filter set (BP 420–480 / BP 495–550 / BP 570–620 / LP 650). Z-stacks were acquired with a 100 nm axial step size, and raw images were reconstructed using the SIM² algorithm in ZEN Blue software (v3.10) to achieve super-resolution datasets.

#### 3. Image processing and quantitative analysis

Z-stacks acquired from lattice structured illumination microscopy (SIM) were processed using SIM² to reconstruct high-resolution images across three fluorescence channels. The resulting datasets were duplicated and preprocessed to facilitate segmentation: Z-projection was generated for each channel and for each plan (sum slices), followed by background subtraction and Gaussian filtering using FIJI [67]. The simplified stacks were then analysed using *Omnipose* [68] to generate segmentation masks for individual bacterial cells. These masks were applied to the original stacks to enable single-cell analysis. This workflow is summarized in Fig. S6. Three-dimensional colocalisation was quantified using Manders’ overlap coefficient, with intensity thresholds determined by Moments method using the JACoP plugin in FIJI [69].

## ACKNOWLEDGEMENTS

N.F.-B and L.B. are grateful to Carmela Giglione, for advice and critical reading of the manuscript, to Philippe Bouloc for support throughout this work and to Patricia Kerboriou for skillful technical assistance. M.B. sincerely thanks Annie Schwartz for her invaluable assistance during the early stages of this project. Funding: This work was supported by the Agence Nationale de la Recherche (grant ANR-15-CE11-0024-01 to E. M., grants ANR-15-CE11-0024-02 and ANR-22-CE44-0017-01 to M.B., and grant ANR-15-CE11-0024-03 to N.F.-B.) and by the Centre National de la Recherche Scientifique. Furthermore, this study benefited from Imagerie-Gif core facility, supported by I’Agence Nationale de la Recherche (FBI ANR-24-INBS-0005 (BIOGEN); SPS ANR-17-EUR-0007, EUR SPS-GSR).

## Supplementary Materials and Methods

### Strain Construction and Characterisation

Strains were all derived from *Salmonella enterica* serovar Typhimurium strain MA3409, a derivative of strain LT2 cured of the Gifsy-1 prophage [1].

#### 1. Construction of a *rho* Merodiploid Strain

To enable unrestricted genetic manipulation of the *rho* gene, we constructed a strain carrying a second chromosomal copy of *rho*. This was achieved in two steps: (*i*) insertion of a *cat* resistance cassette immediately downstream of *rho*, and (*ii*) relocation of the resulting *rho–cat* segment into the *chiP* locus. The choice of the *chiP* locus was initially motivated by the possibility of regulating ectopic *rho* expression via the *chiP*-targeting sRNA ChiX [2]. Accordingly, the *chiP::rho* construct was engineered such that the *rho* open reading frame replaced the *chip* coding sequence, while preserving the native *chip* 5ʹ untranslated region, which contains the ChiX target site. Subsequent analysis revealed, however, that adequate *rho* expression from the *chiP::rho* fusion could be achieved only upon inactivation of the NagC repressor [3], conditions under which ChiX-mediated repression is incomplete due to elevated target mRNA levels. This led us to abandon the use of ChiX as a regulatory device. Nevertheless, the resulting strain, MA11993 (*chiP::rho ΔnagC ΔchiX rho⁺ / pKD46*), was retained and used as the recipient for subsequent genetic manipulations (see next section).

#### 2. Construction of scarless terminal and internal sfGFP fusions to rho in the merodiploid background

Insertions of the sfGFP ORF into the native copy of the *rho* gene in merodiploid strain MA11993 were achieved using two scarless λ Red recombineering strategies. For the N-terminal and internal fusions, a *tetR*-*tetA* cassette was amplified with primers designed to recombine either immediately downstream of rho’s translation start codon or between codons 46 and 49 (the NID position). The resulting PCR fragments were introduced into strain MA11993, yielding tetracycline-resistant, fusaric acid-sensitive strains MA12001 (*rho*2::*tetRA*) and MA12002 (*rho*46::*tetRA*). Next, the sfGFP ORF was PCR-amplified from plasmid pNCM2 [4] with primers designed for recombinational replacement of the *tetRA* cassette. PCR fragments were introduced into strains MA12001 and/or MA12002, and fusaric acid-resistant recombinants were selected at 42°C on Bochner-Maloy plates as described [5]. For the C-terminal sfGFP fusion, a tripartite *tetR*-P^TET^*ccdB*-*kan* cassette (amplified from plasmid pNNB7 [6] with primers AH82 and AH83) was inserted immediately downstream of the native *rho* copy in strain MA11993, generating strain MA14474. This cassette was then replaced by the sfGFP ORF (amplified from plasmid pNCM2 [4] with primers AH84 and AH85). In all cases, multiple clones were picked, single-colony purified, and verified to harbor the desired constructs by colony PCR and Sanger sequencing.

#### 3. Assessing single-copy viability of *rho* terminal and internal sfGFP fusions

A *rho–lacZY* transcriptional fusion was constructed by first introducing an FRT site immediately downstream of *rho* (insertion of an FRT–*kan*–FRT cassette amplified from plasmid pKD4, followed by Flp recombinase–mediated excision of the *kan* element) and then using Flp recombinase to promote integration of plasmid pCE36 at the FRT site [7]. The *rho–lacZ* fusion was transferred into a strain carrying a *cat* gene insertion removing most of the *ilvGEMDA* operon, generating strain MA12221, which is Ilv⁻ and Lac⁺. MA12221 was transduced to prototrophy (Ilv⁺) using P22 lysates prepared from strains harboring the different sfGFP fusions. The appearance of white transductant colonies indicated that the fusion replaced the *rho–lac* region of the recipient and was viable in an unmarked genetic background in the absence of complementation. When lysates from strains carrying either N-terminal (*sfGFP*-*rho*) or C-terminal (*rho*-*sfGFP*) fusions were used, no white colonies were obtained. In contrast, lysates from the strain harboring the intragenic insertion (*rho*::*sfGFP*) yielded approximately 10% white recombinants (Fig. S1). All tested white colonies, upon re-streaking on LB plates, displayed green fluorescence under a 440–460 nm LED light source. These results demonstrate that the intragenic *rho::sfGFP* fusion is compatible with cell viability, whereas both terminal fusions are lethal.

#### 4. Construction of fluorescent protein fusions to *rpoC*

Carboxyl-terminal fusions of *mCherry* and *sfGFP* to the rpoC gene were obtained by λ Red recombineering [4] using DNA fragments amplified from plasmids pROD17 and pNCM2 with primer pairs ppR09—ppQ96 and ppS38—ppS25, respectively (Table S3).

### Protein Purification

Rho::sfGFP and Rho148::mCherry proteins were overexpressed in *E. coli* BL21(DE3)pLysS cells (Novagen) harboring the pET28b-Rho::sfGFP or pET28b-Rho148::mCherry plasmid, respectively. Cells were pelleted and resuspended in Lysis buffer (250 mM NaCl, 1 mM 1-mercaptoethanol, 5 mM EDTA, 0.1 mM DTT, 5% glycerol, 0.05% (w/v) sodium deoxycholate, 50 mM Tris-HCl, pH 7.5) supplemented with cOmplet protease inhibitor cocktail (Roche). Cell lysis was achieved by treatment with Lysozyme (0.5 mg mL^−1^) for 15 min at room temperature followed by sonication using a Bioblock Vibra-Cell 75115 apparatus (40% amplitude, 5 min; 30s on/ 30s off cycles). Insoluble material was eliminated by centrifugation. The bulk of nucleic acids was eliminated by precipitation with 5% (v/v) Polymin-P. Then, proteins were precipitated from the supernatant with 0.5g mL^−1^ ammonium sulfate. After centrifugation, the protein pellet was dissolved in buffer A (100 mM NaCl, 5 mM EDTA, 0.1 mM DTT, 5% glycerol, 10 mM Tris-HCl, pH 7.5) and dialyzed overnight against buffer A. The protein crude extract was loaded onto a Hiload 16/10 SP-sepharose FF column (Cytiva) equilibrated with buffer A and the flow-through (FT) fraction ─ containing the protein of interest ─ was collected. The FT was then directly loaded on a Hiload 16/10 Q-sepharose FF column (Cytiva) equilibrated with buffer A and proteins were eluted from the column with a 0-100% gradient of buffer B (1 M NaCl, 5 mM EDTA, 0.1 mM DTT, 5% glycerol, 10 mM Tris-HCl, pH 7.5). Fractions containing the protein of interest were pooled and loaded on a HiTrap Heparin HP column (Cytiva) equilibrated with buffer A. Proteins were eluted from the column with a 0-100% gradient of buffer B. Fractions of interest were pooled and buffer-exchanged against 2X storage buffer (20 mM Tris-HCl pH 7.9, 5% glycerol, 200 mM KCl, 0.2 mM EDTA), using Amicon Ultra-15 centrifugal units (Millipore). Purified proteins were filtered through 0.22 μm filters, mixed with an equal volume of glycerol, and stored at −20 °C. Protein identity and purity were controlled by high-resolution mass spectrometry at the Mo2ving core facility of CBM (Orléans, France). Concentrations of proteins (expressed in hexamers for Rho derivatives) were determined with the Bio-Rad protein assay (Bio-Rad).

---

**Figure S1.**
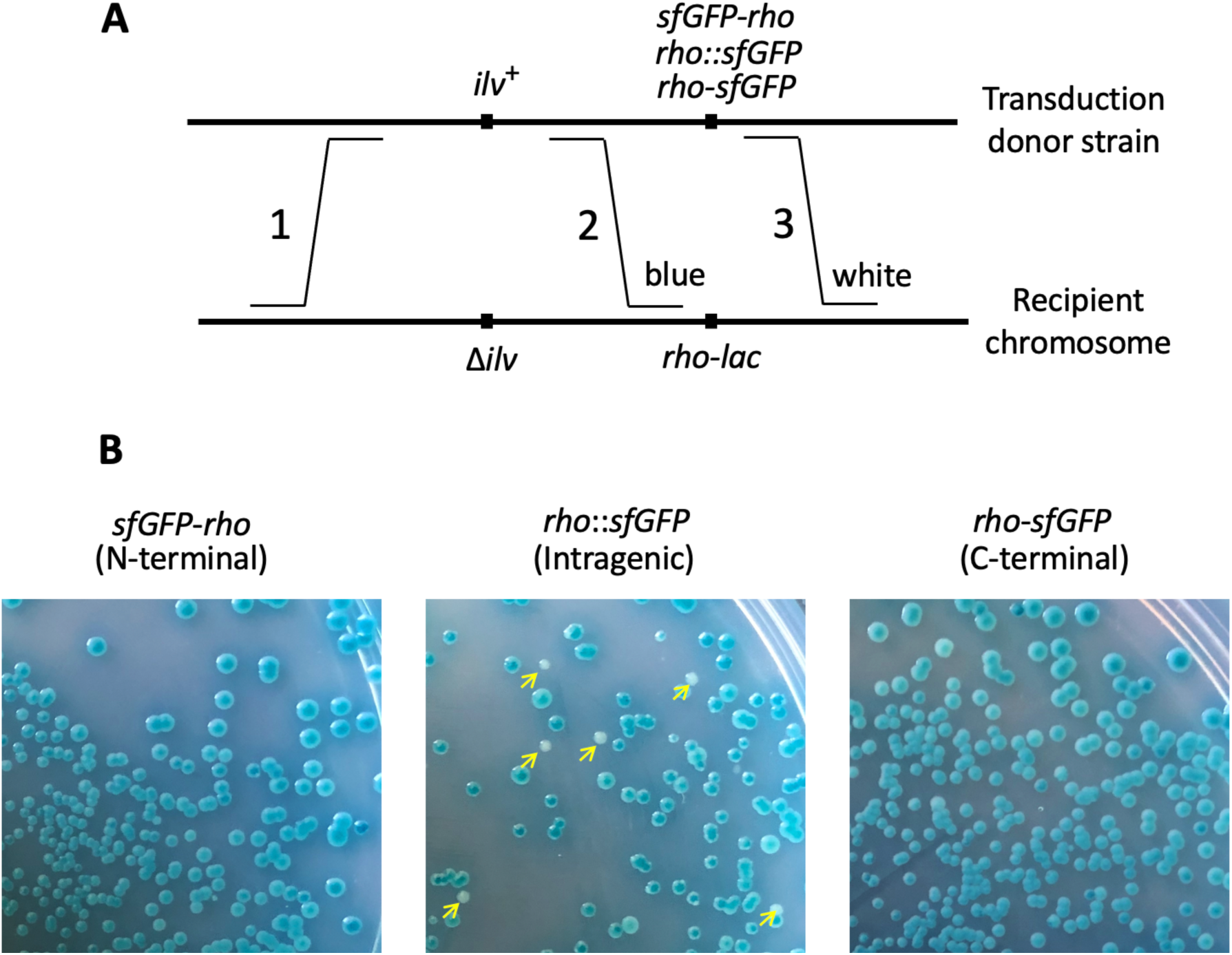
Assaying viability of Rho–sfGFP fusions in single-copy on the chromosome. **A**. Schematic representation of the transductional cross in the *ilv-rho* region. Lysates from strains carrying the three *rho–GFP* fusions were used to infect a Δ*ilv* strain containing a *rho-lac* transcriptional fusion (rho∼lac). Two classes of recombination can produce Ilv⁺ recombinants: crossovers between *ilv* and r*ho* (crosses 1,2; ∼90% frequency) yield blue colonies, whereas crossovers on the opposite side of *rho* (cross 1,3; ∼10% frequency) yield white colonies. **B**. Transduction results. White colonies (yellow arrowheads) are observed only with the *rho*::*sfGFP* donor. Their absence in the other two crosses indicates that the N-and C-terminal fusions (*sfGFP*–*rho* and *rho*–sfGFP) cannot support growth when they are the sole source of Rho activity.

**Figure S2.**
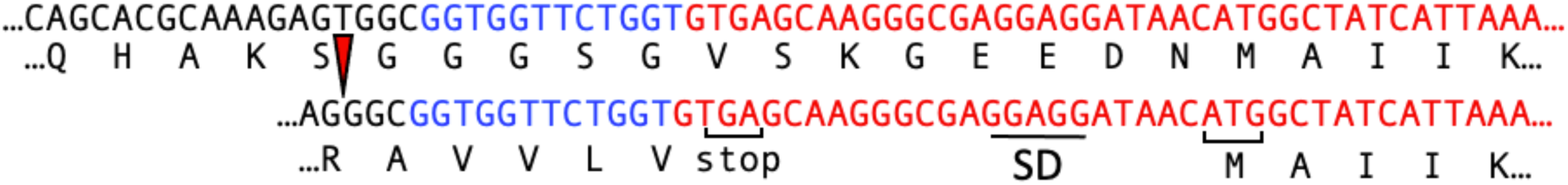
Accidental −1 frameshift mutation at the upstream boundary of the *rho*::*mCherry* insertion reveals a translation re-initiation site at the beginning of the *mCherry* coding sequence.

**Figure S3.**
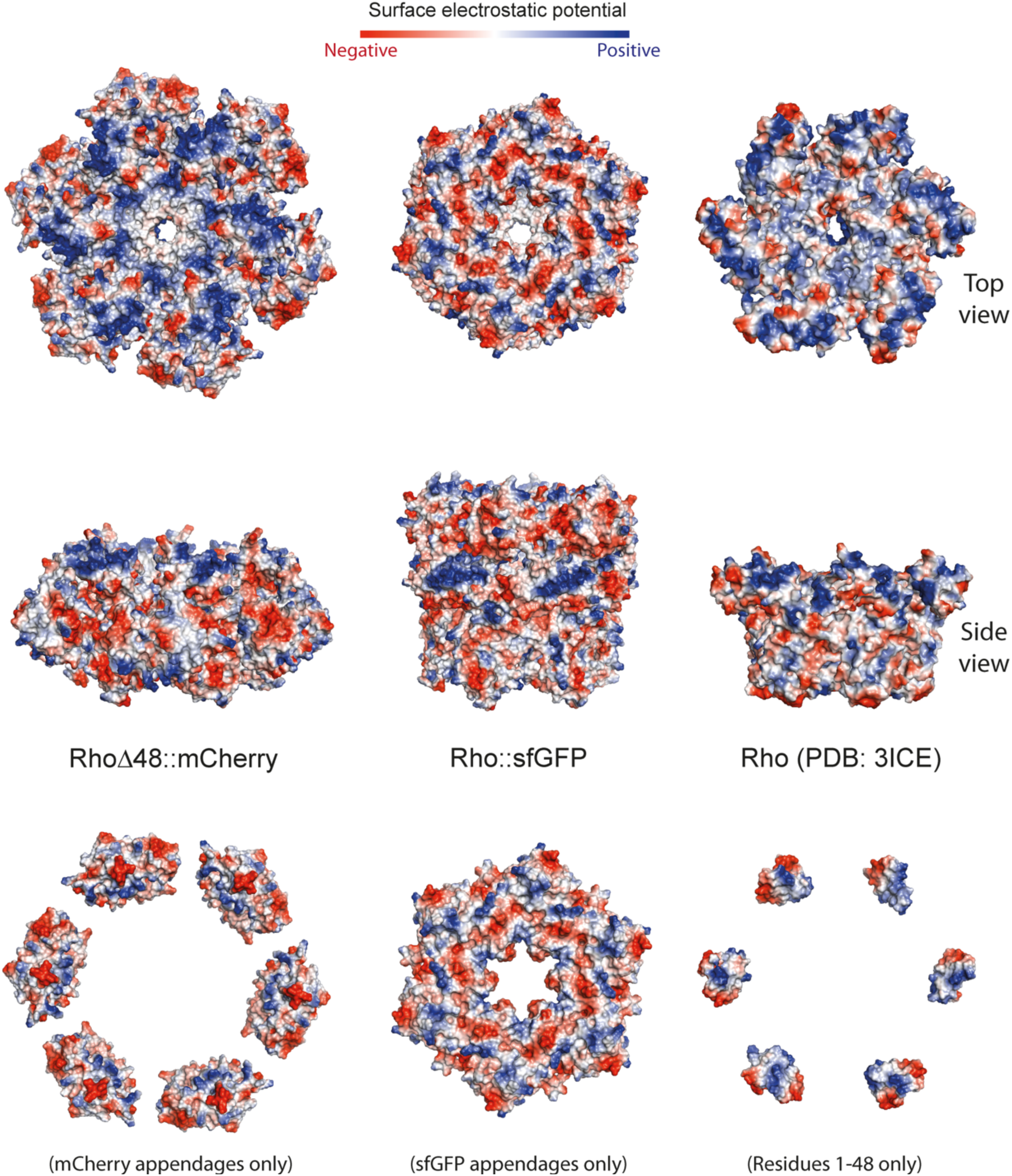
Predicted protein surface electrostatics (Pymol). In WT Rho, the NHB domains and PBS pockets create a crown-like, positively charged surface on the top of the hexamer, facilitating capture of the RNA polyanion. A comparable positive surface is observed atop the RhoΔ48::mCherry model. In contrast, in the Rho::sfGFP model, the top surface potential is predominantly negative. Notably, the positive surface patches on the mCherry appendages are oriented inward, mimicking the positive patches found on the NHB domains (residues 1–48) of WT Rho protomers (bottom).

**Figure S4.**
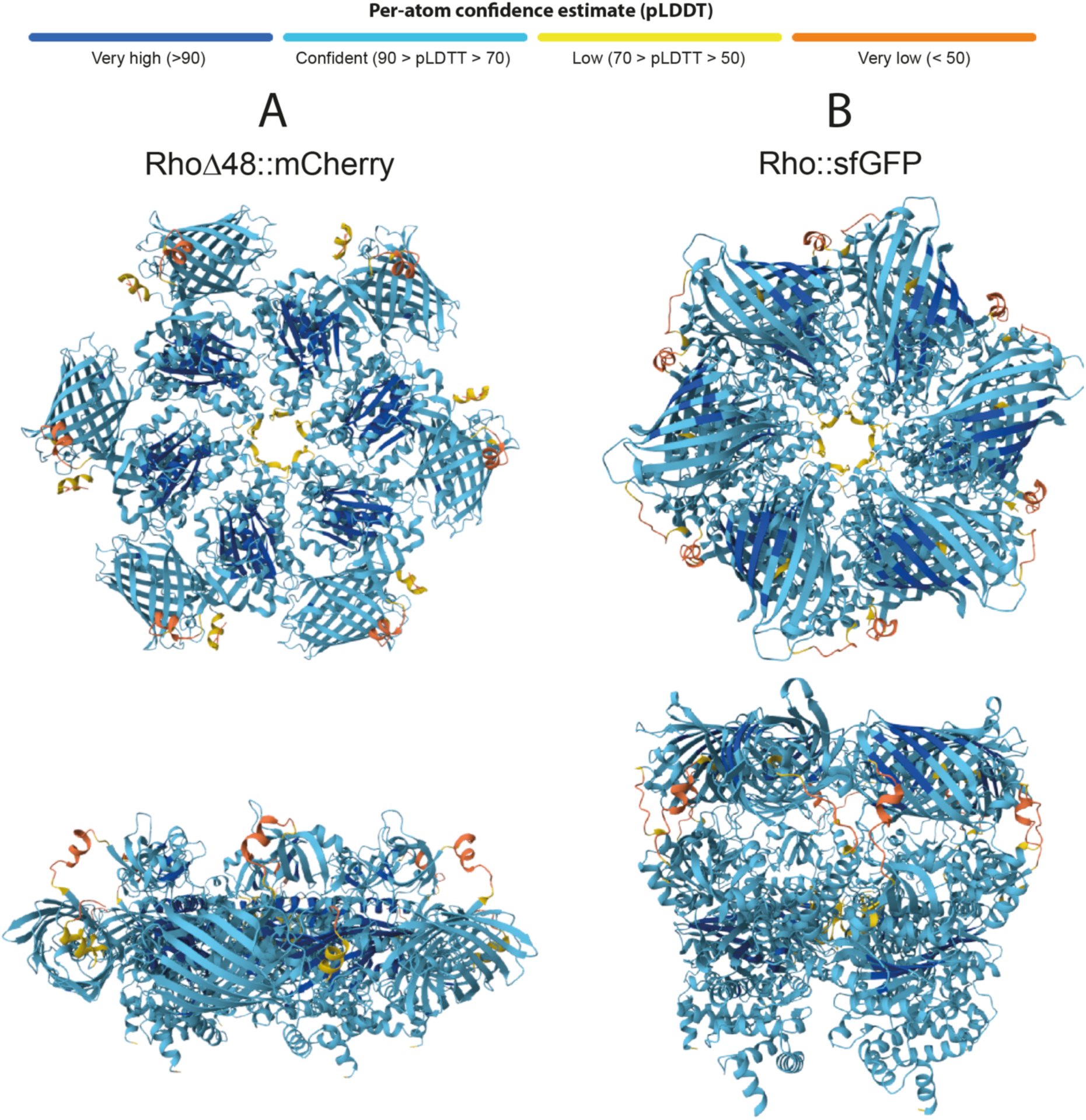
AlphaFold accuracy metrics for RhoΔ48::mCherry and Rho::sfGFP models. The RhoΔ48::mCherry model exhibits a predicted template modeling (pTM) score of 0.51 and an interface predicted template modeling (ipTM) score of 0.47, while the Rho::sfGFP model shows pTM and ipTM scores of 0.48 and 0.45, respectively. Models are colored according to per-atom confidence (pLDDT), with the confidence scale provided at the top of the figure. Both pTM and ipTM scores are used to assess prediction accuracy, with values above 0.5 indicating a predicted fold potentially similar to the native structure.

**Figure S5.**
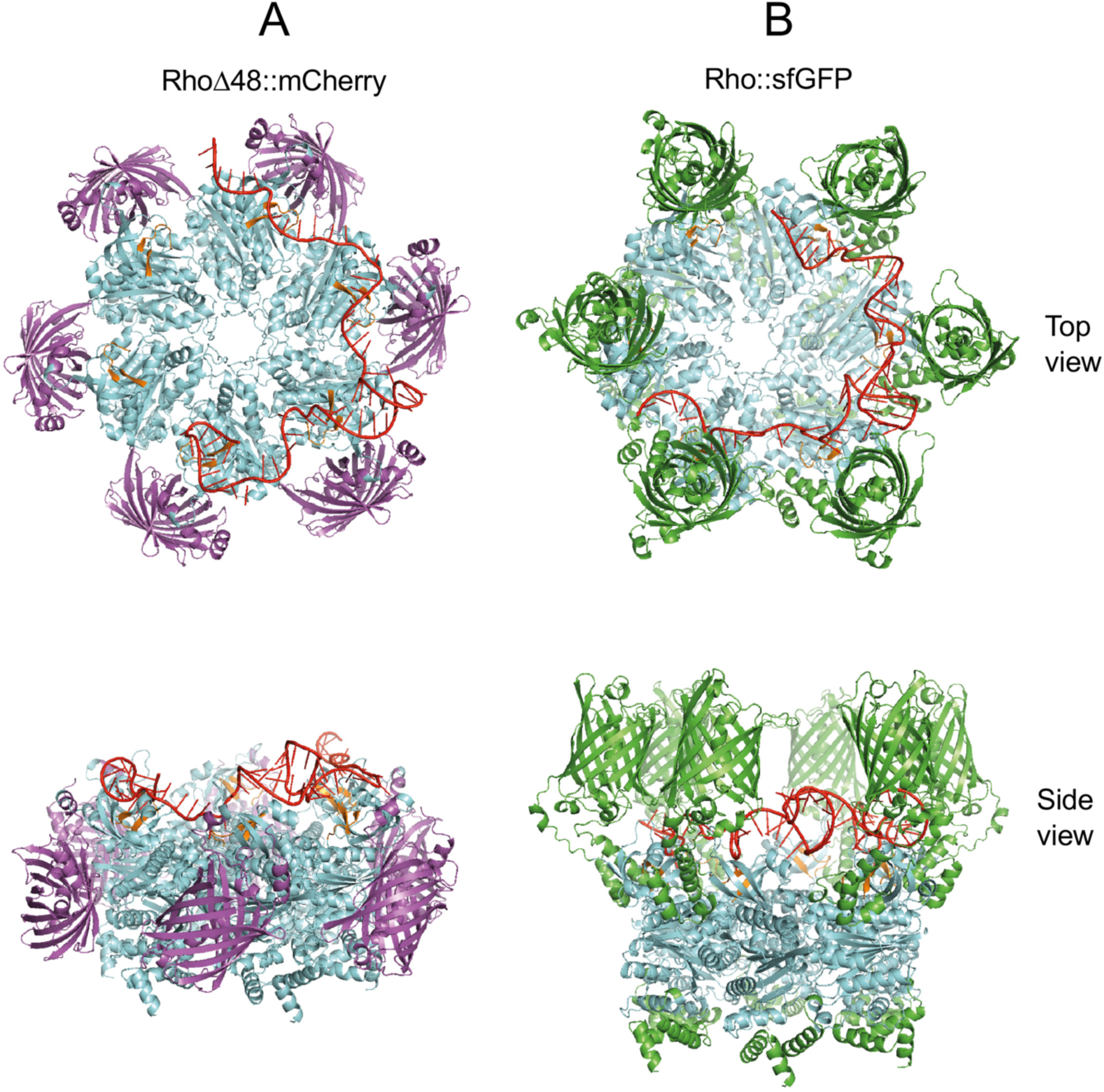
AlphaFold 3 models for the RhoΔ48::mCherry and Rho::sfGFP chimeras in complex with the Rut site RNA of the *λ* tR1 terminator (red). The RNA sequence matches that in a recent cryo-EM structure of the Rho–RNA complex [8]. In the AlphaFold models, the λ tR1 RNA binds as expected to the ring-shaped primary binding site (PBS) of Rho. Comparison with the RNA-less models (Fig. 4) suggests that RNA binding to the Rho::sfGFP chimera requires substandal conformadonal rearrangement ─either through Brownian sampling or acdve displacement by the RNA─ to expose the PBS from the stacked sfGFP appendages.

**Figure S6.**
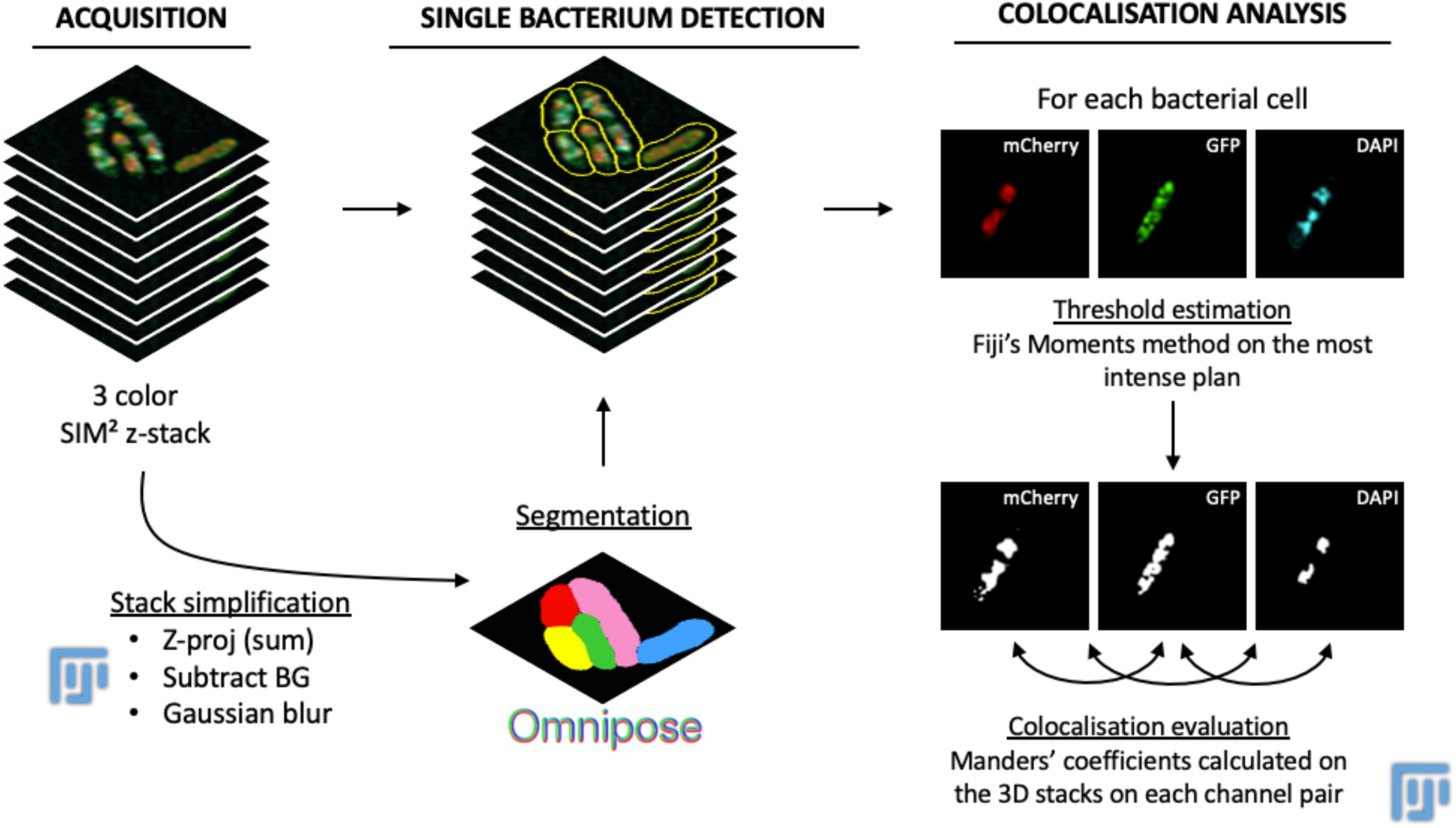
Image processing and quandtadve analysis workflow. Z-stacks acquired from lahce structured illuminadon microscopy (SIM) were processed using SIM² to reconstruct high-resoludon images across three fluorescence channels. The resuldng datasets were duplicated and preprocessed to facilitate segmentadon: Z-projecdon was generated for each channel and for each plan (sum slices), followed by background subtracdon and Gaussian filtering using FIJI. The simplified stacks were then analyzed using *Omnipose* to generate segmentadon masks for individual bacterial cells. These masks were applied to the original stacks to enable single-cell analysis. Three-dimensional colocalisadon was quandfied using Manders’ overlap coefficient, with intensity thresholds determined by Moments method using the JACoP plug-in in FIJI.

**Table S1.**
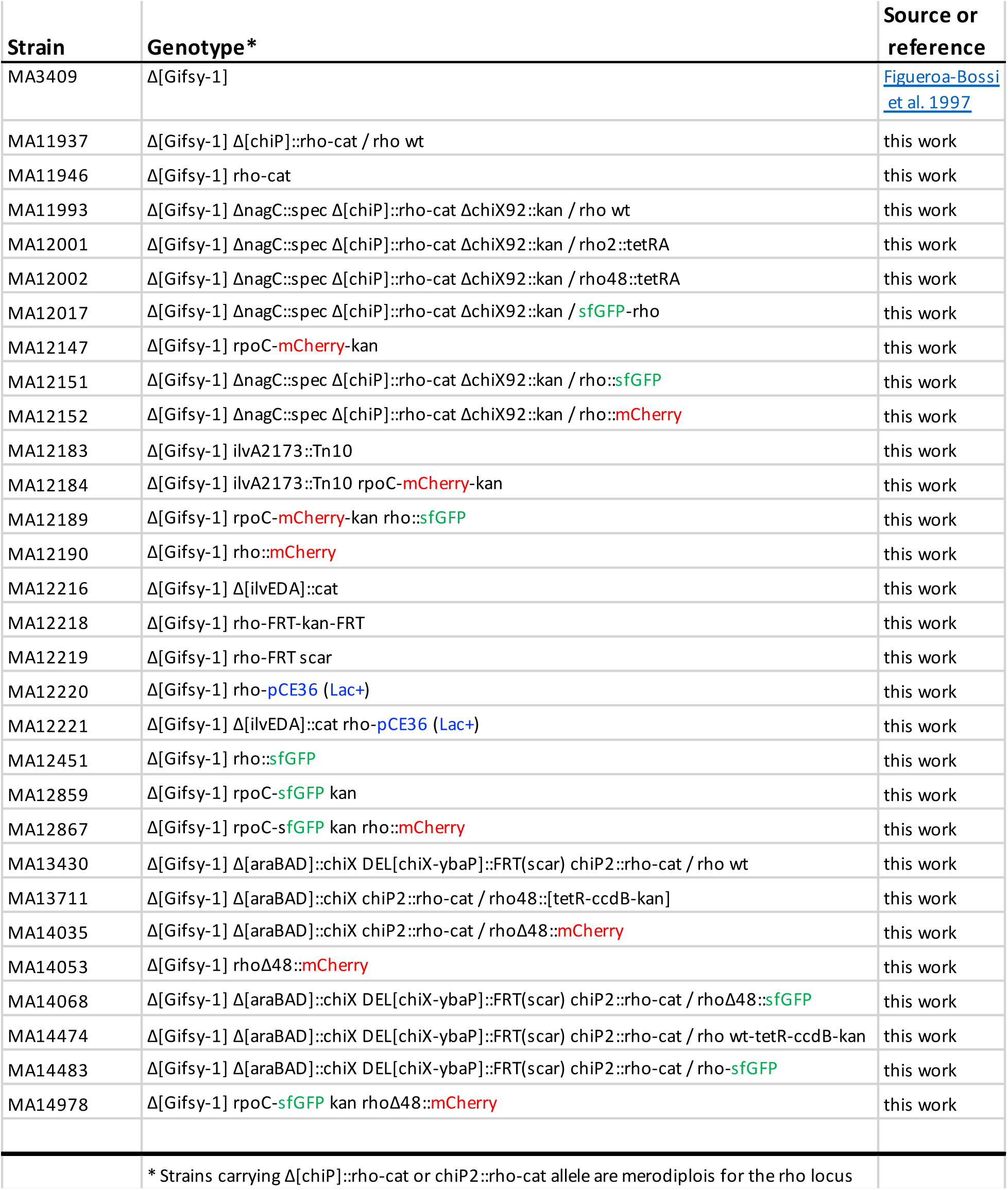
*Salmonella enteric* a serovar Typhimurium strains used in this work.

**Table S2.**
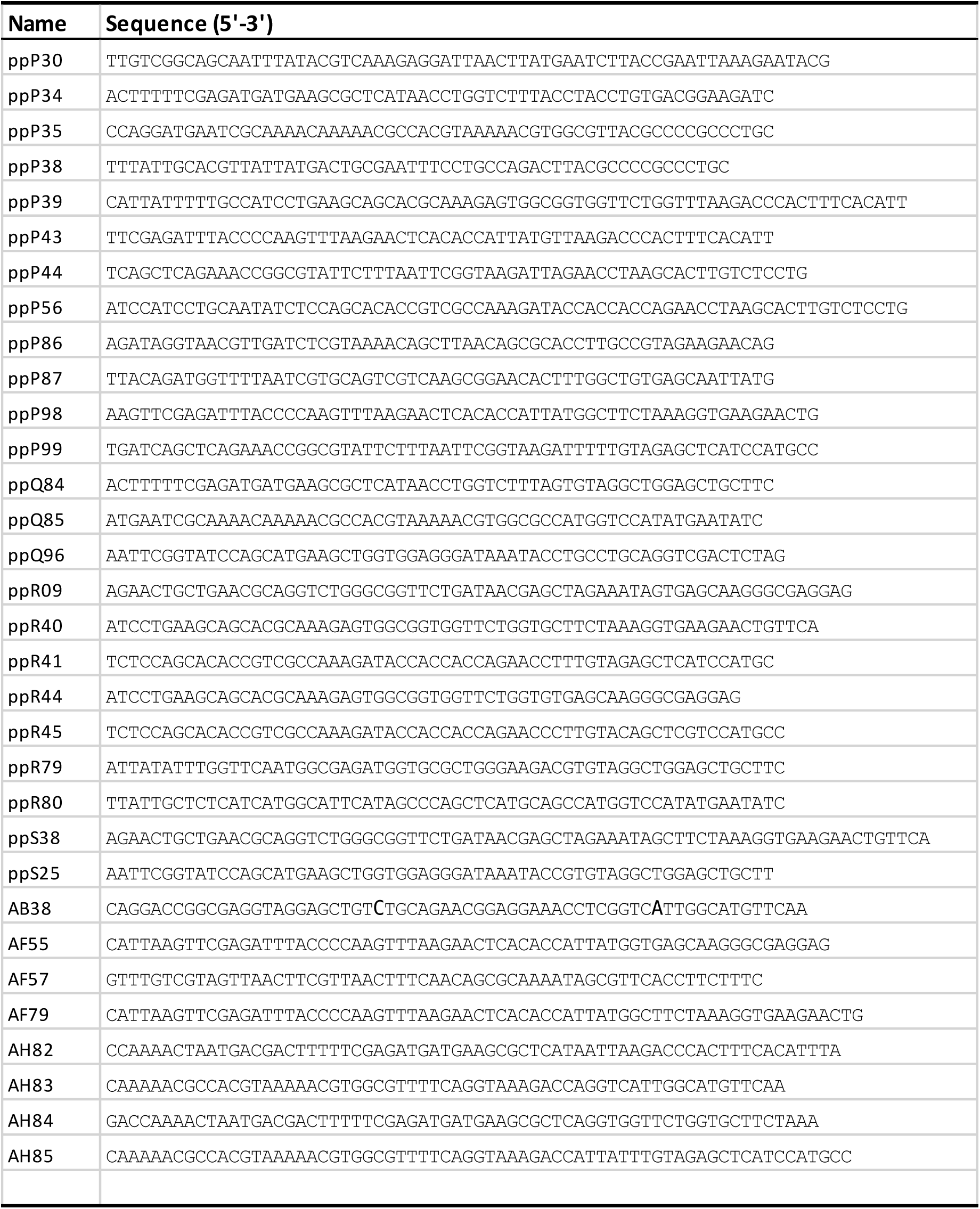
DNA oligonucleotides used for strain construction by λ red recombineering.

**Table S3.** Relevant alleles constructed by λ red recombineering.

| Allele | Primer pair | DNA template | Notes |
| --- | --- | --- | --- |
| rho-cat | ppP34 - ppP35 | plasmid pKD3 | *cat* gene insertion on the 3' side of *rho* gene (used as selectable marked to displace *rho*) |
| ΔchiP::rho-cat | ppP30 - ppP38 | strain MA11946 chromosome | Replacement of *chiP* ORF by *rho* ORF |
| rho2::tetRA | ppP43 - ppP44 | strain MA3397 chromosome | *tetRA* insertion at *rho*'s 2nd codon position |
| rho46::tetRA | ppP39 - ppR56 | strain MA3397 chromosome | *tetRA* insertion at *rho*'s 46th codon position |
| ΔnagC::spec | ppP86 - ppP87 | plasmid pSEB5 | *spec* insertion deleting *nagC* |
| sfGFP-rho | ppP98 - ppP99 | plasmid pNCM2 | N-terminal *sfGFP* fusion to *rho* |
| rho-kan | ppQ84 - ppQ85 | plasmid pKD4 | *kan* gene insertion on the 3' side of *rho* (intermediate step for FLP-mediated insertion of pCE36 plasmid) |
| rpoC-mCherry-ka | ppR09 - ppQ96 | plasmid pROD17 | C-terminal *mCherry* fusion to *rpoC* |
| rho::sfGFP | ppR40 - ppR41 | plasmid pNCM2 | *sfGFP* insertion between *rho* codon positions 46-49 |
| rho::mCherry | ppR44 - ppR45 | plasmid pROD17 | *mCherry* insertion between *rho* codon positions 46-49 |
| $\Delta$[ilvEDA]::cat | ppR79 - ppR80 | plasmid pKD3 | *cat* insertion deleting *ilvEDA* operon |
| rpoC-sfGFP-kan | ppS38 - ppS25 | plasmid pNCM2 | C-terminal *sfGFP* fusion to *rpoC* |
| rhoΔ48::mCherry | AF55 - AF57 | strain MA12190 chromosome | *mCherry* insertion replacing the sequence spanning the initial 48 codons of *rho* |
| rhoΔ48::sfGFP | AF79 - AF57 | strain MA12451 chromosome | *sfGFP* insertion replacing the sequence spanning the initial 48 codons of *rho* |
| rho tetR-ccdB-ka | AH82 - AH83 | plasmid pNNB7 | Counter-selectable *tetR-ccdB-kan* cassette immediately downstream from *rho* |
| rho-sfGFP | AH84 - AH85 | plasmid pNCM2 | C-terminal *sfGFP* fusion to *rho* |

**Table S4.** Colocalization analyses, raw data (continues on next page)

| **MA12189 (rho::sfGFP/ RpoC-mCherry) EXPONENTIAL** |  |  |  |  |  |  |
| --- | --- | --- | --- | --- | --- | --- |
| **Rho:RpoC / RpoC:Rho** |  |  |  |  |  |  |
| Image N. | 1 | 2 | 3 | 4 |  |  |
| Rho centroids colocalizing with RpoC centroids | 42 | 52 | 61 | 83 |  |  |
| Total Rho centroids | 196 | 233 | 247 | 417 |  |  |
| % colocalization Rho:RpoC | 21,4285714 | 22,3175966 | 24,6963563 | 19,9040767 |  |  |
| RpoC centroids colocalizing with Rho centroids | 42 | 52 | 61 | 83 |  |  |
| Total RpoC centroids | 281 | 313 | 280 | 436 |  |  |
| % colocalization RpoC:Rho | 14,9466192 | 16,6134185 | 21,7857143 | 19,0366972 |  |  |
| **Rho:DNA / DNA:Rho** |  |  |  |  |  |  |
| Image N. | 1 | 2 | 3 | 4 |  |  |
| Rho centroids colocalizing with DNA centroids | 24 | 26 | 35 | 53 |  |  |
| Total Rho centroids | 196 | 233 | 247 | 417 |  |  |
| % colocalization Rho:DNA | 12,244898 | 11,1587983 | 14,1700405 | 12,7098321 |  |  |
| DNA centroids colocalizing with Rho centroids | 24 | 26 | 35 | 53 |  |  |
| Total DNA centroids | 213 | 255 | 213 | 358 |  |  |
| % colocalization DNA:Rho | 11,2676056 | 10,1960784 | 16,4319249 | 14,8044693 |  |  |
| **RpoC:DNA / DNA:RpoC** |  |  |  |  |  |  |
| Image N. | 1 | 2 | 3 | 4 |  |  |
| RpoC centroids colocalizing with DNA centroids | 62 | 100 | 100 | 83 |  |  |
| Total RpoC centroids | 281 | 313 | 280 | 436 |  |  |
| % colocalization RpoC:DNA | 22,0640569 | 31,9488818 | 35,7142857 | 19,0366972 |  |  |
| DNA centroids colocalizing with RpoC centroids | 62 | 100 | 100 | 83 |  |  |
| Total DNA centroids | 213 | 258 | 213 | 358 |  |  |
| % colocalization DNA:Rho | 29,1079812 | 38,7596899 | 46,9483568 | 23,1843575 |  |  |
| **MA12189 (rho::sfGFP/ RpoC-mCherry) STATIONARY** |  |  |  |  |  |  |
| **Rho:RpoC / RpoC:Rho** |  |  |  |  |  |  |
| Image N. | 1 | 2 | 3 | 4 | 5 | 6 |
| Rho centroids colocalizing with RpoC centroids | 38 | 32 | 44 | 31 | 38 | 35 |
| Total Rho centroids | 245 | 253 | 234 | 195 | 245 | 252 |
| % colocalization Rho:RpoC | 15,5102041 | 12,6482213 | 18,8034188 | 15,8974359 | 15,5102041 | 13,8888889 |
| RpoC centroids colocalizing with Rho centroids | 38 | 32 | 44 | 31 | 38 | 35 |
| Total RpoC centroids | 161 | 168 | 163 | 154 | 178 | 186 |
| % colocalization RpoC:Rho | 23,6024845 | 19,047619 | 26,993865 | 20,1298701 | 21,3483146 | 18,8172043 |
| **Rho:DNA / DNA:Rho** |  |  |  |  |  |  |
| Image N. | 1 | 2 | 3 | 4 | 5 | 6 |
| Rho centroids colocalizing with DNA centroids | 28 | 23 | 29 | 24 | 25 | 29 |
| Total Rho centroids | 245 | 253 | 234 | 195 | 245 | 252 |
| % colocalization Rho:DNA | 11,4285714 | 9,09090909 | 12,3931624 | 12,3076923 | 10,2040816 | 11,5079365 |
| DNA centroids colocalizing with Rho centroids | 28 | 23 | 29 | 24 | 25 | 29 |
| Total DNA centroids | 147 | 170 | 155 | 149 | 165 | 182 |
| % colocalization DNA:Rho | 19,047619 | 13,5294118 | 18,7096774 | 16,1073826 | 15,1515152 | 15,9340659 |
| **RpoC:DNA / DNA:RpoC** |  |  |  |  |  |  |
| Image N. | 1 | 2 | 3 | 4 | 5 | 6 |
| RpoC centroids colocalizing with DNA centroids | 60 | 77 | 63 | 60 | 76 | 85 |
| Total RpoC centroids | 161 | 168 | 163 | 154 | 178 | 186 |
| % colocalization RpoC:DNA | 37,2670807 | 45,8333333 | 38,6503067 | 38,961039 | 42,6966292 | 45,6989247 |
| DNA centroids colocalizing with RpoC centroids | 60 | 77 | 63 | 60 | 76 | 85 |
| Total DNA centroids | 147 | 170 | 155 | 149 | 165 | 182 |
| % colocalization DNA:Rho | 40,8163265 | 45,2941176 | 40,6451613 | 40,2684564 | 46,0606061 | 46,7032967 |

**Table S4. Colocalization analyses, raw data (continued from previous page)**
| **MA14978 (RhoΔ48::mCherry / RpoC-sfGFP) EXPONENTIAL** |  |  |  |  |  |  |
| --- | --- | --- | --- | --- | --- | --- |
| **Rho:RpoC / RpoC:Rho** |  |  |  |  |  |  |
| Image N. | 1 | 2 | 3 | 4 | 5 | 6 |
| Rho centroids colocalizing with RpoC centroids | 32 | 30 | 16 | 26 | 30 | 75 |
| Total Rho centroids | 509 | 265 | 248 | 260 | 248 | 462 |
| % colocalization Rho:RpoC | 6,28683694 | 11,3207547 | 6,4516129 | 10 | 12,0967742 | 16,2337662 |
| RpoC centroids colocalizing with Rho centroids | 32 | 30 | 16 | 26 | 30 | 75 |
| Total RpoC centroids | 297 | 166 | 197 | 201 | 164 | 425 |
| % colocalization RpoC:Rho | 10,7744108 | 18,0722892 | 8,12182741 | 12,9353234 | 18,2926829 | 17,6470588 |
| **Rho:DNA / DNA:Rho** |  |  |  |  |  |  |
| Image N. | 1 | 2 | 3 | 4 | 5 | 6 |
| Rho centroids colocalizing with DNA centroids | 38 | 39 | 27 | 28 | 31 | 89 |
| Total Rho centroids | 509 | 265 | 248 | 260 | 248 | 462 |
| % colocalization Rho:DNA | 7,46561886 | 14,7169811 | 10,8870968 | 10,7692308 | 12,5 | 19,2640693 |
| DNA centroids colocalizing with Rho centroids | 38 | 39 | 27 | 28 | 31 | 89 |
| Total DNA centroids | 241 | 130 | 144 | 170 | 129 | 362 |
| % colocalization DNA:Rho | 15,7676349 | 30 | 18,75 | 16,4705882 | 24,0310078 | 24,5856354 |
| **RpoC:DNA / DNA:RpoC** |  |  |  |  |  |  |
| Image N. | 1 | 2 | 3 | 4 | 5 | 6 |
| RpoC centroids colocalizing with DNA centroids | 64 | 89 | 79 | 114 | 81 | 251 |
| Total RpoC centroids | 297 | 166 | 197 | 201 | 164 | 425 |
| % colocalization RpoC:DNA | 21,5488215 | 53,6144578 | 40,1015228 | 56,7164179 | 49,3902439 | 59,0588235 |
| DNA centroids colocalizing with RpoC centroids | 64 | 89 | 79 | 114 | 81 | 251 |
| Total DNA centroids | 241 | 130 | 144 | 170 | 129 | 362 |
| % colocalization DNA:Rho | 26,5560166 | 68,4615385 | 54,8611111 | 67,0588235 | 62,7906977 | 69,3370166 |
| **MA14978 (RhoΔ48::mCherry / RpoC-sfGFP) STATIONARY** |  |  |  |  |  |  |
| **Rho:RpoC / RpoC:Rho** |  |  |  |  |  |  |
| Image N. | 1 | 2 | 3 | 4 | 5 | 6 |
| Rho centroids colocalizing with RpoC centroids | 60 | 52 | 73 | 58 | 54 | 67 |
| Total Rho centroids | 232 | 260 | 283 | 214 | 240 | 254 |
| % colocalization Rho:RpoC | 25,862069 | 20 | 25,795053 | 27,1028037 | 22,5 | 26,3779528 |
| RpoC centroids colocalizing with Rho centroids | 60 | 52 | 73 | 58 | 54 | 67 |
| Total RpoC centroids | 183 | 194 | 230 | 171 | 211 | 196 |
| % colocalization RpoC:Rho | 32,7868852 | 26,8041237 | 31,7391304 | 33,9181287 | 25,5924171 | 34,1836735 |
| **Rho:DNA / DNA:Rho** |  |  |  |  |  |  |
| Image N. | 1 | 2 | 3 | 4 | 5 | 6 |
| Rho centroids colocalizing with DNA centroids | 28 | 38 | 35 | 32 | 33 | 45 |
| Total Rho centroids | 232 | 260 | 283 | 214 | 240 | 254 |
| % colocalization Rho:DNA | 12,0689655 | 14,6153846 | 12,3674912 | 14,953271 | 13,75 | 17,7165354 |
| DNA centroids colocalizing with Rho centroids | 28 | 38 | 35 | 32 | 33 | 45 |
| Total DNA centroids | 187 | 191 | 237 | 173 | 217 | 198 |
| % colocalization DNA:Rho | 14,973262 | 19,895288 | 14,7679325 | 18,4971098 | 15,2073733 | 22,7272727 |
| **RpoC:DNA / DNA:RpoC** |  |  |  |  |  |  |
| Image N. | 1 | 2 | 3 | 4 | 5 | 6 |
| RpoC centroids colocalizing with DNA centroids | 66 | 99 | 130 | 96 | 107 | 110 |
| Total RpoC centroids | 183 | 194 | 230 | 171 | 211 | 196 |
| % colocalization RpoC:DNA | 36,0655738 | 51,0309278 | 56,5217391 | 56,1403509 | 50,7109005 | 56,122449 |
| DNA centroids colocalizing with RpoC centroids | 66 | 99 | 130 | 96 | 107 | 110 |
| Total DNA centroids | 187 | 191 | 237 | 173 | 217 | 198 |
| % colocalization DNA:Rho | 35,2941176 | 51,8324607 | 54,8523207 | 55,4913295 | 49,3087558 | 55,5555556 |

**Table S5.**
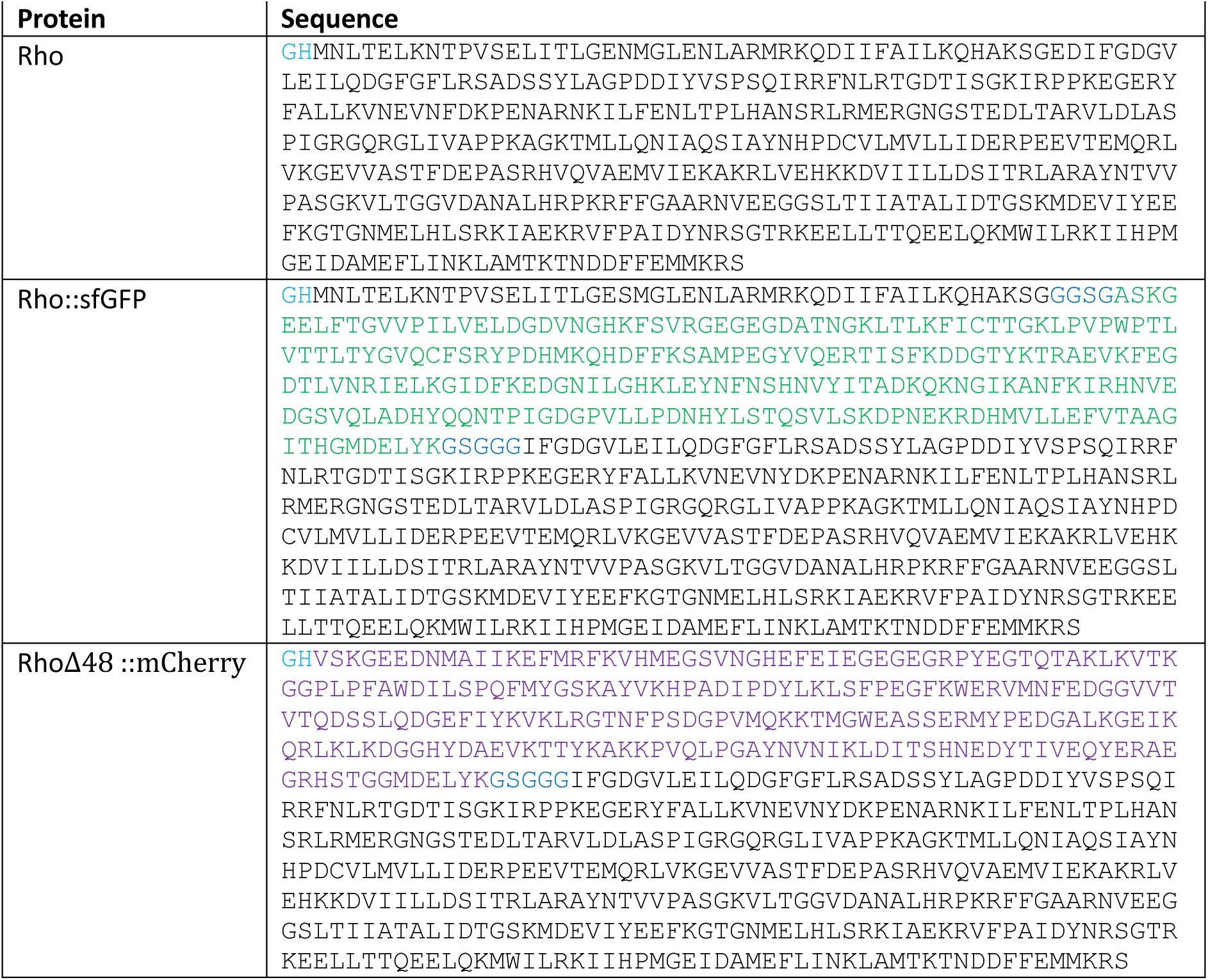
Sequences of Rho protein derivatives used in *in vitro* assays.

## References

1. Ray-Soni, A., M.J. Bellecourt, and R. Landick, Mechanisms of Bacterial Transcription Termination: All Good Things Must End. Annu Rev Biochem, 2016. 85: p. 319–47.

2. Bidnenko, V., et al., Termination factor Rho: From the control of pervasive transcription to cell fate determination in Bacillus subtilis. PLoS Genet, 2017. 13(7): p. e1006909.

3. Botella, L., et al., Depleting Mycobacterium tuberculosis of the transcription termination factor Rho causes pervasive transcription and rapid death. Nat Commun, 2017. 8: p. 14731.

4. Figueroa-Bossi, N., et al., Pervasive transcription enhances the accessibility of H-NS-silenced promoters and generates bistability in Salmonella virulence gene expression. Proc Natl Acad Sci U S A, 2022. 119(30): p. e2203011119.

5. Peters, J.M., et al., Rho and NusG suppress pervasive antisense transcription in Escherichia coli. Genes Dev, 2012. 26(23): p. 2621–33.

6. Do, T.D., et al., Rho-dependent transcription termination: mechanisms and roles in bacterial fitness and adaptation to environmental changes. RNA, 2025. 31(9): p. 1207–1234.

7. Hollands, K., A. Sevostiyanova, and E.A. Groisman, Unusually long-lived pause required for regulation of a Rho-dependent transcription terminator. Proc Natl Acad Sci U S A, 2014. 111(19): p. E1999–2007.

8. Molodtsov, V., et al., Structural basis of Rho-dependent transcription termination. Nature, 2023. 614(7947): p. 367–374.

9. Martinez, A., C.M. Burns, and J.P. Richardson, Residues in the RNP1-like sequence motif of Rho protein are involved in RNA-binding affinity and discrimination. J Mol Biol, 1996. 257(5): p. 909–18.

10. Shashni, R., et al., Suppression of in vivo Rho-dependent transcription termination defects: evidence for kinetically controlled steps. Microbiology (Reading), 2012. 158(Pt 6): p. 1468–1481.

11. Steinmetz, E.J. and T. Platt, Evidence supporting a tethered tracking model for helicase activity of Escherichia coli Rho factor. Proc Natl Acad Sci U S A, 1994. 91(4): p. 1401–5.

12. Epshtein, V., et al., An allosteric mechanism of Rho-dependent transcription termination. Nature, 2010. 463(7278): p. 245–9.

13. Hao, Z., et al., Pre-termination Transcription Complex: Structure and Function. Mol Cell, 2021. 81(2): p. 281–292 e8.

14. Said, N., et al., Steps toward translocation-independent RNA polymerase inactivation by terminator ATPase rho. Science, 2021. 371(6524).

15. Song, E., et al., Rho-dependent transcription termination proceeds via three routes. Nat Commun, 2022. 13(1): p. 1663.

16. D’Heygere, F., M. Rabhi, and M. Boudvillain, Phyletic distribution and conservation of the bacterial transcription termination factor Rho. Microbiology, 2013. 159(Pt 7): p. 1423–36.

17. Moreira, S.M., et al., Diversification of the Rho transcription termination factor in bacteria. Nucleic Acids Res, 2024. 52(15): p. 8979–8997.

18. Krypotou, E., et al., Bacteria require phase separation for fitness in the mammalian gut. Science, 2023. 379(6637): p. 1149–1156.

19. Warren Norris, M.A.H., D.M. Plaskon, and R. Tamayo, Phase Variation of Flagella and Toxins in Clostridioides difficile is Mediated by Selective Rho-dependent Termination. J Mol Biol, 2024. 436(6): p. 168456.

20. Pallares, I., V. Iglesias, and S. Ventura, The Rho Termination Factor of Clostridium botulinum Contains a Prion-Like Domain with a Highly Amyloidogenic Core. Front Microbiol, 2015. 6: p. 1516.

21. Yuan, A.H. and A. Hochschild, A bacterial global regulator forms a prion. Science, 2017. 355(6321): p. 198–201.

22. Gjorgjevikj, D., et al., The Psu protein of phage satellite P4 inhibits transcription termination factor rho by forced hyper-oligomerization. Nat Commun, 2025. 16(1): p. 550.

23. Wang, B., et al., Nucleotide-induced hyper-oligomerization inactivates transcription termination factor rho. Nat Commun, 2025. 16(1): p. 1653.

24. Barik, S., P. Bhattacharya, and A. Das, Autogenous regulation of transcription termination factor Rho. J Mol Biol, 1985. 182(4): p. 495–508.

25. Matsumoto, Y., et al., Autogenous regulation of the gene for transcription termination factor rho in Escherichia coli: localisation and function of its attenuators. J Bacteriol, 1986. 166(3): p. 945–58.

26. Ladouceur, A.M., et al., Clusters of bacterial RNA polymerase are biomolecular condensates that assemble through liquid-liquid phase separation. Proc Natl Acad Sci U S A, 2020. 117(31): p. 18540–18549.

27. Balbontín, R., N. Figueroa-Bossi, and L. Bossi, Construction of Single-Copy Fluorescent Protein Fusions by One-Step Recombineering. Cold Spring Harb Protoc, 2023. 2023(9): p. 651–662.

28. Anderson, P. and J. Roth, Spontaneous tandem genetic duplications in Salmonella typhimurium arise by unequal recombination between rRNA (rrn) cistrons. Proc Natl Acad Sci U S A, 1981. 78(5): p. 3113–7.

29. Fages-Lartaud, M., et al., mCherry contains a fluorescent protein isoform that interferes with its reporter function. Front Bioeng Biotechnol, 2022. 10: p. 892138.

30. Zou, L.L. and J.P. Richardson, Enhancement of transcription termination factor rho activity with potassium glutamate. J Biol Chem, 1991. 266(16): p. 10201–9.

31. Lowery, C. and J.P. Richardson, Characterization of the nucleoside triphosphate phosphohydrolase (ATPase) activity of RNA synthesis termination factor p. II. Influence of synthetic RNA homopolymers and random copolymers on the reaction. J Biol Chem, 1977. 252(4): p. 1381–5.

32. Abramson, J., et al., Accurate structure prediction of biomolecular interactions with AlphaFold 3. Nature, 2024. 630(8016): p. 493–500.

33. Taghbalout, A., Q. Yang, and V. Arluison, The Escherichia coli RNA processing and degradation machinery is compartmentalized within an organized cellular network. Biochem J, 2014. 458(1): p. 11–22.

34. Jain, S., A. Behera, and R. Sen, DNA binding of an RNA helicase bacterial transcription terminator. Biochem J, 2025. 482(3): p. 103–117.

35. Zhu, Y., M. Mustafi, and J.C. Weisshaar, Biophysical Properties of Escherichia coli Cytoplasm in Stationary Phase by Superresolution Fluorescence Microscopy. mBio, 2020. 11(3).

36. Cabrera, J.E. and D.J. Jin, The distribution of RNA polymerase in Escherichia coli is dynamic and sensitive to environmental cues. Mol Microbiol, 2003. 50(5): p. 1493–505.

37. Endesfelder, U., et al., Multiscale spatial organization of RNA polymerase in Escherichia coli. Biophys J, 2013. 105(1): p. 172–81.

38. Fan, J., et al., RNA polymerase redistribution supports growth in E. coli strains with a minimal number of rRNA operons. Nucleic Acids Res, 2023. 51(15): p. 8085–8101.

39. Gaal, T., et al., Colocalisation of distant chromosomal loci in space in E. coli: a bacterial nucleolus. Genes Dev, 2016. 30(20): p. 2272–2285.

40. Lewis, P.J., S.D. Thaker, and J. Errington, Compartmentalization of transcription and translation in Bacillus subtilis. EMBO J, 2000. 19(4): p. 710–8.

41. Stracy, M., et al., Live-cell superresolution microscopy reveals the organization of RNA polymerase in the bacterial nucleoid. Proc Natl Acad Sci U S A, 2015. 112(32): p. E4390–9.

42. Weng, X., et al., Spatial organization of RNA polymerase and its relationship with transcription in Escherichia coli. Proc Natl Acad Sci U S A, 2019. 116(40): p. 20115–20123.

43. Morgan, E.A., Antitermination mechanisms in rRNA operons of Escherichia coli. J Bacteriol, 1986. 168(1): p. 1–5.

44. Nudler, E. and M.E. Gottesman, Transcription termination and anti-termination in E. coli. Genes Cells, 2002. 7(8): p. 755–68.

45. Schmidt, A., et al., The quantitative and condition-dependent Escherichia coli proteome. Nat Biotechnol, 2016. 34(1): p. 104–10.

46. Ditlev, J.A., L.B. Case, and M.K. Rosen, Who’s In and Who’s Out-Compositional Control of Biomolecular Condensates. J Mol Biol, 2018. 430(23): p. 4666–4684.

47. Al-Husini, N., et al., alpha-Proteobacterial RNA Degradosomes Assemble Liquid-Liquid Phase-Separated RNP Bodies. Mol Cell, 2018. 71(6): p. 1027–1039 e14.

48. Montero Llopis, P., et al., Spatial organization of the flow of genetic information in bacteria. Nature, 2010. 466(7302): p. 77–81.

49. Jäger, S., et al., An mRNA degrading complex in Rhodobacter capsulatus. Nucleic Acids Res, 2001. 29(22): p. 4581–8.

50. Jäger, S., et al., Composition and activity of the Rhodobacter capsulatus degradosome vary under different oxygen concentrations. J Mol Microbiol Biotechnol, 2004. 7(3): p. 148–54.

51. Tuckerman, J.R., G. Gonzalez, and M.A. Gilles-Gonzalez, Cyclic di-GMP activation of polynucleotide phosphorylase signal-dependent RNA processing. J Mol Biol, 2011. 407(5): p. 633–9.

52. Liou, G.G., et al., RNA degradosomes exist in vivo in Escherichia coli as multicomponent complexes associated with the cytoplasmic membrane via the N-terminal region of ribonuclease E. Proc Natl Acad Sci U S A, 2001. 98(1): p. 63–8.

53. Taghbalout, A. and L. Rothfield, RNaseE and the other constituents of the RNA degradosome are components of the bacterial cytoskeleton. Proc Natl Acad Sci U S A, 2007. 104(5): p. 1667–72.

54. Lilleengen, K., Typing of Salmonella typhimurium by means of bacteriophage. Acta Pathol. Microbiol. Scand., 1948. 77(Suppl): p. 2–125.

55. Figueroa-Bossi, N., et al., Unsuspected prophage-like elements in Salmonella typhimurium. Mol Microbiol, 1997. 25(1): p. 161–73.

56. Bertani, G., Lysogeny at mid-twentieth century: P1, P2, and other experimental systems. J Bacteriol, 2004. 186(3): p. 595–600.

57. Maloy, S.R. and J.R. Roth, Regulation of proline utilization in Salmonella typhimurium: characterization of put::Mu d(Ap, lac) operon fusions. J Bacteriol, 1983. 154(2): p. 561–8.

58. Schmieger, H., Phage P22-mutants with increased or decreased transduction abilities. Mol Gen Genet, 1972. 119(1): p. 75–88.

59. Datsenko, K.A. and B.L. Wanner, One-step inactivation of chromosomal genes in Escherichia coli K-12 using PCR products. Proc Natl Acad Sci U S A, 2000. 97(12): p. 6640–5.

60. Figueroa-Bossi, N. and L. Bossi, Recombineering applications for the mutational analysis of bacterial RNA-binding proteins and their sites of action. Methods Mol Biol, 2015. 1259: p. 103–16.

61. Boudvillain, M., et al., Simple enzymatic assays for the in vitro motor activity of transcription termination factor Rho from Escherichia coli. Methods Mol Biol, 2010. 587: p. 137–54.

62. Simon, I., et al., A Large Insertion Domain in the Rho Factor From a Low G + C, Gram-negative Bacterium is Critical for RNA Binding and Transcription Termination Activity. J Mol Biol, 2021. 433(15): p. 167060.

63. Delaleau, M., et al., Rho-dependent transcriptional switches regulate the bacterial response to cold shock. Mol Cell, 2024. 84(18): p. 3482–3496 e7.

64. Rabhi, M., et al., Mutagenesis-based evidence for an asymmetric configuration of the ring-shaped transcription termination factor Rho. J Mol Biol, 2011. 405(2): p. 497–518.

65. Lau, L.F., J.W. Roberts, and R. Wu, Transcription terminates at lambda tR1 in three clusters. Proc Natl Acad Sci U S A, 1982. 79(20): p. 6171–5.

66. Walmacq, C., A.R. Rahmouni, and M. Boudvillain, Influence of substrate composition on the helicase activity of transcription termination factor Rho: reduced processivity of Rho hexamers during unwinding of RNA-DNA hybrid regions. J Mol Biol, 2004. 342(2): p. 403–20.

67. Schindelin, J., et al., Fiji: an open-source platform for biological-image analysis. Nat Methods, 2012. 9(7): p. 676–82.

68. Cutler, K.J., et al., Omnipose: a high-precision morphology-independent solution for bacterial cell segmentation. Nat Methods, 2022. 19(11): p. 1438–1448.

69. Bolte, S. and F.P. Cordelieres, A guided tour into subcellular colocalisation analysis in light microscopy. J Microsc, 2006. 224(Pt 3): p. 213–32.

## References (Supplementary Methods)

1. Figueroa-Bossi, N., et al., Unsuspected prophage-like elements in Salmonella typhimurium. Mol Microbiol, 1997. 25(1): p. 161–73.

2. Figueroa-Bossi, N., et al., Caught at its own game: regulatory small RNA inactivated by an inducible transcript mimicking its target. Genes Dev, 2009. 23(17): p. 2004–15.

3. Plumbridge, J., et al., Interplay of transcriptional and small RNA-dependent control mechanisms regulates chitosugar uptake in Escherichia coli and Salmonella. Mol Microbiol, 2014. 92(4): p. 648–58.

4. Balbonvn, R., N. Figueroa-Bossi, and L. Bossi, Construction of Single-Copy Fluorescent Protein Fusions by One-Step Recombineering. Cold Spring Harb Protoc, 2023. 2023(9): p. 651–662.

5. Figueroa-Bossi, N. and L. Bossi, Recombineering applications for the mutational analysis of bacterial RNA-binding proteins and their sites of action. Methods Mol Biol, 2015. 1259: p. 103–16.

6. Figueroa-Bossi, N., R. Balbonwn, and L. Bossi, Scarless DNA Recombineering. Cold Spring Harb Protoc, 2023. 2023(9): p. pdb prot107857.

7. Ellermeier, C.D., A. Janakiraman, and J.M. Slauch, Construction of targeted single copy lac fusions using lambda Red and FLP-mediated site-specific recombination in bacteria. Gene, 2002. 290(1-2): p. 153–61.

8. Molodtsov, V., et al., Structural basis of Rho-dependent transcription termination. Nature, 2023. 614(7947): p. 367–374.

